# Age, sex, and cell type-resolved hypothalamic gene expression across the pubertal transition in mice

**DOI:** 10.1101/2023.10.12.562121

**Authors:** Dustin J. Sokolowski, Huayun Hou, Kyoko E. Yuki, Anna Roy, Cadia Chan, Mariela Faykoo-Martinez, Matt Hudson, Liis Uusküla-Reimand, Anna Goldenberg, Mark R. Palmert, Michael D. Wilson

## Abstract

Although the hypothalamus plays a critical role in the regulation of puberty, more research is needed to identify the gene regulatory networks that control pubertal timing. Here, we investigate the age-, sex- and cell-type-specific gene regulation in the hypothalamus across the pubertal transition. We used RNA-seq to profile hypothalamic gene expression in male and female mice at five time points spanning the onset of puberty (postnatal days (PD) 12, 22, 27, 32, and 37). By combining this data with hypothalamic scRNA-seq data of pre- and post-pubertal mice, we were able to assign gene expression changes to their cell types of origin. In our colony, pubertal onset occurs earlier in male mice allowing us to focus on genes whose expression is dynamic across ages and offset between sexes and to explore bases of sex effects. Our age-by-sex pattern of expression enriched for biological pathways involved hormone production, neuronal activation, and glial maturation. Additionally, we found a dramatic expansion of oligodendrocytes precursor cells into mature oligodendrocytes spanning the pre-pubertal (PD12) to peri-pubertal (PD27) timepoints, and that genes driving this expansion enrich for genes involved in pubertal regulation. Together, by incorporating multiple biological timepoints with male and female mice simultaneously, our work furthers the understanding of gene and cell-type changes that accompany the development of secondary sex characteristics in both sexes.

## Introduction

Puberty is a fundamental period of mammalian development when individuals reach sexual maturity and can produce gametes. Despite being a nearly universal event, pubertal timing within the population varies and is known to be influenced by genetic and environmental factors [1, 2], though much of its variation remains unexplained. Rare mutations in several genes lead to pubertal disorders such as central precocious puberty (CPP), defined as abnormally early pubertal initiation, and hypogonadotropic hypogonadism (HH), defined as delayed or absent puberty, due to misregulated or missing gonadotropic hormones [3, 4]. Recent genome-wide association studies (GWAS) investigating the age of menarche in females and age of voice breaking in males [1, 5] have identified common variants related to pubertal timing, which influence the timing of puberty in the general population and are associated with important health outcomes. Specifically, early puberty is associated with increased risk of later life health outcomes of such as cancer, diabetes, and cardiovascular disease, while late puberty is associated increased risk of osteoporosis and mental health disorders [1, 5–7]. Furthermore, environmental factors such as diet, body mass index (BMI), prenatal growth, and psychosocial experience are associated with differences in pubertal timing [7, 8].

Puberty is initiated in the hypothalamus by pulses of gonadotropin-releasing hormone (GnRH) that then stimulate the pituitary gland to increase secretion of luteinizing hormone (LH) and follicle stimulating hormone (FSH) increases in frequency and amplitude. This cascade begins an organism-wide feedback loop involving many genes, cell types, and gene regulatory mechanisms [1, 9, 10]. Previous studies have investigated hypothalamic regulation during puberty and have discovered a growing list of gene-regulatory mechanisms that can directly regulate pubertal timing [2, 11–13]. These groundbreaking studies include the epigenetic mechanisms activating and inhibiting pubertal onset and spatial transcriptomic programs associated with post-natal development in the female rat hypothalamus [9, 12, 14–16].

Puberty is an inherently sex-biased process as it results in the development of secondary sex characteristics, and males and females undergo pubertal timing at different ages. In humans, puberty occurs earlier in females, and in rodent models sex-differences are also seen, with male mice in our colony undergoing earlier development than females [10]. While the sex-specific physiological differences between males and females are different, many of the same gene regulatory mechanisms are likely present, but playing slightly different roles [5, 17, 18]. For example, in humans, the same variant in the *LIN28B* gene is associated with puberty-relevant phenotypes in both males and females [18]. However, mouse models of *Lin28b* and *Lin28a* knockouts revealed sexually dimorphic phenotypes related to body weight and pubertal development [17]. Accordingly, measuring genome-wide pubertal dynamics in the hypothalamus while accounting for multiple timepoints, sex, and cell types should yield greater insight into pubertal development and disease [2]. From the perspective of measuring hypothalamic gene expression across pubertal development, genes whose developmental trajectories are offset between male and female mice provide powerful candidates for differential pubertal regulation, making age and sex pertinent variables to study pubertal regulation.

Only a few studies have characterized hypothalamic gene expression across the pubertal transition, and fewer have incorporated both age and sex into their design [10, 19]. Our lab previously utilized multiplexed qPCR to measure the expression of 178 candidate puberty GWAS and disease-related genes at PD12, 22, 27, 32, and 37 in many tissues, including the hypothalamus [10]. There, we found that most age-biased expression in the hypothalamus occurred before puberty, perhaps reflecting the cellular development occurring before puberty [2, 16, 20]. Our lab has also performed RNA-seq through 3’UTR profiling of these same mice in the pituitary gland, where we discovered cell-type specific expression and cellular trajectories that became increasingly sexually dimorphic through the pubertal transition [19]. Recently, Han et al. interrogated the premammillary nucleus and arcuate nucleus transcriptomes of female mice between postnatal day (PD) 20 and diestrus females (PD50-PD60), as well as a leptin-inducible model of puberty in adult mice, highlighting the importance of neuropil and somatodendritic organization [20]. These authors focused on how leptin may lead to puberty-relevant transcriptomic changes rather than investigating pubertal development over time or across sexes [20].

The above findings are consistent with recognition that cellular complexity of the hypothalamus plays an important role in its ability to regulate many biological processes. Accordingly, researchers have employed single-cell transcriptomics to profile the complexity of the hypothalamus at multiple pre-natal and post-natal developmental timepoints [16, 21–23]. In the mouse, Kim et al., 2020 used scRNA-seq to study embryonic and early postnatal hypothalamic development [23]. They included two timepoints surrounding puberty, PD14 and PD45, as developmental endpoints, which could be re-analyzed to study puberty [23]. Importantly, these scRNA-seq data can be integrated with bulk RNA-seq dataset to incorporate cell-type specificity into the study while maintaining multiple biological replicates across different ages and sexes in a single experimental batch [24–26].

Thus, in this study, we measured hypothalamic gene expression in male and female mice at five timepoints spanning pubertal transition. We identified age-biased genes primarily associated with cellular development and a smaller set involved in hormonal regulation. We also identified a key subset of genes whose expression is conditional on age and sex, but the observed sex effects were not robust or plentiful, consistent with our previous work suggesting that the pituitary may have larger role in sex differences in pubertal timing than the hypothalamus. Using hypothalamic scRNA-seq (as above), we mapped the age and sex conditional genes to their most likely cell type of origin, including neurons and oligodendrocytes. We further integrated these data to discover that substantial oligodendrocyte expansion occurs before and during puberty in mice, which is interesting in that many identified genes associated with oligodendrocyte expansion have been previously implicated in modulation of pubertal timing in humans. Lastly, the gene expression distribution of all genes can be freely interacted with using our Shiny App (<u>wilsonlab-sickkids-uoft.shinyapps.io/hypothalamus_gene_shiny/</u>), so that researchers can identify the natural gene expression patterns of their genes of interest before perturbation. Overall, by analyzing the hypothalamic transcriptome during pubertal development in a manner that simultaneously incorporates age and sex, we identified novel genes involved in cellular composition and hormonal regulation in the pubertal hypothalamus.

## Materials and Methods

### Animal and tissue collection

Tissue dissection and RNA extraction follow the protocol in Hou et al., 2017 as the same samples were utilized [10]. We collected the hypothami of 48 C57BL/6 mice at PD12, 22, 27, 32, and 37 in males and females (4-5 mice per age/sex).

### Library preparation and sequencing

RNA-seq libraries were prepared using an automated - QuantSeq 3’mRNA-seq (Lexogen GmbH, Vienna) and Agilent NGS Workstation (Agilent Technologies, Santa Clara) at The Centre for Applied Genomics (TCAG) (Toronto, Canada) as per the manufacturer’s protocol (UTRSeq). The automated QuantSeq 3’mRNA-seq library construction was described in detail in Hou et al., 2022 [19]. Briefly, 250 ng of total RNA spiked-in with ERCC Spike-In Control Mix 1 (Ambion) as per the manufacturer’s protocol was used to generate cDNA. cDNA was amplified with 17 PCR cycles as determined by qPCR analysis using the PCR Add-on kit (Lexogen). The resulting libraries were quantified with Qubit DNA High Sensitivity assay (ThermoFisher). Fragment sizes were analyzed on the Agilent Bioanalyzer using the High Sensitivity DNA assay prior to sequencing. Single-read 50-bp sequencing was performed at TCAG on an Illumina HiSeq2500 Rapid Run or V4 flowcell (Illumina, San Diego) with cycles extended to 68 bp.

### Read processing

Reads from technical replicates were merged prior to downstream analyses. Fastqc (http://www.bioinformatics.babraham.ac.uk/projects/fastqc/) was used to examine the quality of sequenced reads. A customized script to trim both the polyAs and adapters at the end of the reads [19] was used. The script implemented a “back search” strategy to account for cases where a mixture of adapters and polyAs were seen at the end of the reads. In addition, the first 12 nucleotides were trimmed with Cutadapt [27] based on the manufacturer’s recommendations. Only reads longer than 36 bp after trimming were used for future analyses. After trimming, Fastqc was performed again to examine read quality, and over-represented reads, namely reads mapping to *BC1* (brain cytoplasmic 1), were removed. Trimmed and filtered reads were aligned to the genome using a splice-aware aligner, STAR (version 2.5.1b), with default settings except “--outFilterMismatchNoverLmax 0.05” for QuantSeq [28]. Quality control (QC) of mapped RNA-seq reads was performed using Qualimap version 2.2.1 (Supplementary Table S1). Read signal was visualized with the UCSC genome browser [29, 30]. Reads were assigned to genes using featureCounts (version 1.5.3) [31] with parameters “ -s 1 -Q 255 -t exon -O”. Gene models were obtained from GENCODE M11 [32, 33].

### Count processing and evaluation

Counts successfully aligned to GENCODE M11 [33] were normalized based on ERCC spike-ins using RUVseq [34]. Genes with fewer than 5 reads were removed before upper-quartile normalization was completed with the betweenLaneNormalization() function [34]. Finally, ERCC spike-ins were used to normalize counts using the RUVg(), yielding the final normalized count matrix [34]. All samples were correlated to one another using Pearson’s correlation of all genes before being plotted with the ComplexHeatmap package [35]. Genes overlapping the RNA-seq and qPCR data of the same samples [10] were correlated using Pearson’s correlation analysis. Principal component analysis (PCA) of samples was performed with the “prcomp()” function [36] before being plotted with ggplot2 [37].

### Differential expression analysis

Pairwise differential gene expression analysis was completed across ages and sexes. Differentially expressed genes were calculated using the DESeq2 R package [38]. Genes were considered differentially expressed if they had a false discovery rate (FDR)-adjusted p-value < 0.05 and an absolute-value fold-change 1.5. Sex comparisons were completed at each timepoint, while age comparisons within each sex were completed between days 12 and 22, 22 and 27, 27 and 32, and 32 and 37.

### Varimax rotation principal component analysis

Principal component analysis (PCA) is a dimensionality reduction technique used to reduce every individual mouse’s global gene expression pattern into a smaller set of orthogonal vectors [36] (N_components_ = Nmice = 48). Varimax rotation decreases the distance between PCs and mice by adjusting the PC axes such that samples will more closely align with one varimax rotated PC (vrPC) [39]. By leveraging vrPC scores, defined by the location of a sample of a PC axis, we identified which vrPCs are associated with age, sex, and an age-by-sex interaction by completing a two-way ANOVA of timepoint and sex on vrPC scores. By leveraging vrPC loadings, defined by the association between a gene and PC, we measured which genes are represented by individual vrPCs.

Normalized count data and PC scores were used to generate varimax-rotated PCs with the “varimax” function in R [39, 40]. Varimax-rotated PC loadings and scores were acquired using pracma [41]. A loading is a gene’s coefficient to the vrPC, while the score is a sample’s coefficient to a rotated vrPC [39, 40]. The association between scores, age, and sex was measured using two-way ANOVA. Multiple-test correction using the FDR was applied using the p.adjust() function in R [42]. The FDRs of the vrPCs with an associated main effect or interaction were plotted with ggplot2 [37]. We designated that genes with loading greater than three standard deviations from the mean loading are associated with a vrPC. We picked three standard deviations by inspecting a qqplot of loadings with the qqnorm function. Genes associated with a vrPC were re-ordered by the loading magnitude for downstream analysis.

### Pathway and human RNA-seq enrichment analysis

Pathway enrichment of fused gene lists (e.g., PD12 vs. PD22, males and females) was completed using the ActivePathways R package [43]. Briefly, ActivePathways takes the p-values from different related gene lists (e.g., PD12 vs. PD22 - males, PD12 vs. PD22 - females) and fuses them using Brown’s extension of Fisher’s method [43]. Then, it computes pathway enrichment of each individual gene list and the fused gene list using a p-value-ranked Hypergeometric test. The resulting statistics provide pathway enrichments annotated to each DEG list and their integrated p-values [43]. We used the “Mouse_GO_AllPathways_no_GO_iea_September_01_2022_symbol.gmt” gene set database from (http://download.baderlab.org/EM_Genesets/), which systematically curates a gene set list from multiple sources (Gene Ontology, Reactome, Panther, etc.) as our pathway enrichment database [44].

Pathway enrichment for gene lists without p-values following a multivariate normal distribution (i.e., vrPC-associated genes, oligodendrocyte-pseudotime associated genes) was completed using the g:ProfileR R package using an FDR correction, with genes detected in the RNA-seq dataset as the custom background and with GO:BP, GO:MF, and GO:CC being queried [44, 45]. Here, “genes” represent oligodendrocyte-pseudotime associated genes or associated loadings ordered by FDR-adjusted p-value or vrPC loading, respectively. Biological pathways identified by integrating developmental changes across sexes were completed with ActivePathways [43].

We used the Differential Expression Enrichment Tool (DEET) to compare our age-biased DEGs and vrPC-associated genes to 3162 consistently reprocessed sets of DEGs derived from The Cancer Genome Atlas (TCGA), Genotype-Tissue Expression Consortium (GTEx), and from various studies within the Sequencing Read Archive (SRA) [46–50]. To test the human RNA-seq DEG-set enrichment of fused gene lists (i.e., PD12 vs. PD22 male, PD12 vs. PD22 female) We extracted their “DEET_gmt_DE” object, which stores their DEG sets as a generic pathway-enrichment database. We used this gene set as the pathway enrichment database using ActivePathways, using the same parameters as in our fused pathway enrichment. Next, we ran the “DEET_enrich()” function to measure which of our age-biased DEG lists and vrPC- associated genes enriched for publicly available human DEG sets. DEET_enrich() also identifies DEG comparisons whose overlapping DEGS also has a correlated fold-change, suggesting that the shared DEGs and pathways may be under shared regulation [44]. Correlation plots were generated using the “DEET_enrichment_plot” with default parameters. Lastly, we enriched our neuron-neuroendocrine mapping age-by-sex associated genes with LepRb+ cells in the hypothalamus by overlapping Trap-seq+ genes from Alison et al. [51] and testing for over-representation with a Fisher’s exact test. Pathway enrichment plots for ActivePathways, traditional pathway enrichment, and DEET, were completed using the “DEET_enrichment_plot” and “process_and_plot_DEET_enrich” functions.

### Processing of public hypothalamic scRNA-seq data

Filtered gene-barcode matrix files for the PD14 and two PD45 samples were downloaded from the Gene Expression Omnibus (GEO) series GSE132355 (P14: GSM3860745, P45-rep1: GSM3860746, P45-rep2: GSM3860747) [23].

Counts were processed and integrated using the “process_dgTmatrix_lists” function in scMappR [24], including all genes and scTransform [52] as options. Briefly, “process_dgTmatrix_lists” is a wrapper for Seurat V4 and scTransform [52, 53] before cell-type labeling with cell-type enrichment of the CellMarker [54] and Panglao [55] databases. In our preprocessing of these data, we used the Integration Anchors with Canonical Correlation Analysis, a rigorous recommended batch correction method [52, 56] because the PD14 mice were from the CD1 strain and the PD45 mice were from the C57BL/6J mice. While we may have lost some developmental signal through this rigorous batch correction [57], the major cell-type markers and developmental trajectories we observed would be more reliable and translatable to our bulk RNA-seq.

Cells were first labeled with the cell types provided by the original authors [23]. we further applied the cluster labels and cell-type markers generated from “process_dgTmatrix_lists” [24] to provide further specificity to these cell-types. For example, “oligodendrocytes” contained clusters “4”, “24”, “17” and “24” which could be annotated to “oligodendrocyte precursors”, “developing oligodendrocytes”, and “mature oligodendrocytes”. Cells with a different major cell-type label (i.e., neuron vs. glia) between the original author and this analysis and cell-types whose markers were primarily mitochondrial genes were discarded for differential, proportion, and trajectory analyses.

### Age-biased cell type-specific gene expression and cell-type proportion in scRNA-seq data

Age-biased cell-type proportion changes were measured with Fisher’s exact-test [58]. Age-biased genes within each cell type were measured using the Model-Based Analysis of Single-cell Transcriptomics (MAST) within the “FindMarkers” wrapper in Seurat [53, 59]. We filtered genes with an FDR-adjusted p-value < 0.05 and required the gene to be expressed in >25% of the cells in either age group.

### Cell-type deconvolution

All defined cell types and all samples were used in cell-type deconvolution analysis. We completed RNA-seq deconvolution with DeconRNA-seq [60], Digital Cell Quantification (DCQ) [61], Whole Gene Correlation Network Analysis (WGCNA) [62], Cibersort and CibersortX [63, 64], and Cell population mapping (CPM) [65], MuSiC R package [26], and BayesPrism [25]. For all methods, the bulk RNA-seq dataset were the same RUV-seq normalized counts [34] and the scRNA-seq data were the SCTransform-normalized counts [52]. Cell-type proportions from the MuSic and MuSiC-NNLS methods were computed simultaneously with the “music_prop” function, using default parameters [26]. We then correlated the predicted cell-type proportion at PND12 with the cell-type proportions of scRNA-seq data at PND14 and the predicted cell-type proportion at PND37 to the cell-type proportions of scRNA-seq data at PND45. We used the tool with the highest correlation to the scRNA-seq data, Music-NNLS, for downstream analysis [26]. For all downstream analyses, we estimated cell-type proportions with scRNA-seq PD14 and PD45 timepoints combined. We used DeconRNA-seq to calculate cwFold-changes in scMappR because it had the strongest correlation between predicted cell-type proportions and scRNA-seq cell-type proportions of the three allowed RNA-seq deconvolution methods for the scMappR tool, namely DeconRNA-seq, WGCNA, and DCQ [24, 60–62, 66]. We then used the cell-type proportions estimated by MuSiC-NNLS to assign genes to cell types because the cell-type proportion filter of gene–cell-type assignment can use any deconvolution method [24, 26, 60]. The association between cell-type proportion, sex, and age was measured with two-way ANOVA.

### Cell-type specificity of bulk differentially expressed genes

We used scMappR [24] to generate a signature matrix from the scRNA-seq data in Kim et al., 2020 [23] by using the “generes_to_heatmap” function in conjunction with our previously labeled cell types. We then calculated cell-weighted Fold-Changes (cwFCs) for genes associated with varimax-rotated PC 16 with the “scMappR_and_pathway_analysis” function before sorting each DEG into the cell type driving it with the “cwFoldChange_evaluate” function [24].

We next used scMappR to assign genes to their cell type of origin based on the differential expression of the genes between the conditions of interest in a specific cell-type. To calculate the cell-weighted fold-change statistic, we inputted the bulk fold-change from PD12 and PD32, as these timepoints have the largest distance in the varimax-rotated PC 16.

### Cell-type trajectories of scRNA-seq data

Cell-type trajectories were measured in oligodendrocytes (oligodendrocyte precursors [OPCs], developing oligodendrocytes [DOs], and mature oligodendrocytes [MOs]) using the “slingshot” R package [67] with default parameters other than setting the “extension” parameter to “n”. The starting cell type in each trajectory was set as the most “PD14-biased” cluster, namely the “Oligodendrocyte precursor”. We analyzed which genes had expression patterns associated with pseudotime trajectories using the “tradeSeq” [68] R package. We used the minimum number of allowable knots from the “evaluateK” function to fit the negative-binomial generalized additive model with the “fitGAM” function [68]. Then, we tested the association between genes and trajectories with the “associationTest” function [68], and corrected p-values with the “fdr” correction. Genes with an FDR-adjusted p-value < 0.05 and a fold-change > 1.5 remained for downstream analysis. We used the “predictSmooth” [68] function paired with the scale function to generate columns for heatmaps. We identified genes based on their overlap with mouse TFs from ENCODE [69], puberty GWAS genes [1], and genes associated with varimax-rotated PC 16. We plotted the expression of these genes along pseudotime with the Pheatmap function.

## Results

### Global transcriptomic view of the postnatal mouse hypothalamus across the pubertal transition in males and females

To track the transcriptomic dynamics of the postnatal mouse hypothalamus, we measured genome-wide gene expression using 3’UTR profiling in male and female C57BL/6J mouse hypothalamus samples collected at five post-natal days (PDs) corresponding to early development (day 12), pre-pubertal (day 22), pubertal (day 27 in males, and day 32 in females), and post-pubertal (day 37) stages (N = 4-5 per sex/age) (Figure 1A). Specially, we applied an automated, high-throughput RNA-seq platform to complete 3’UTR sequencing of all mice in a single experimental batch.

**Figure 1.**
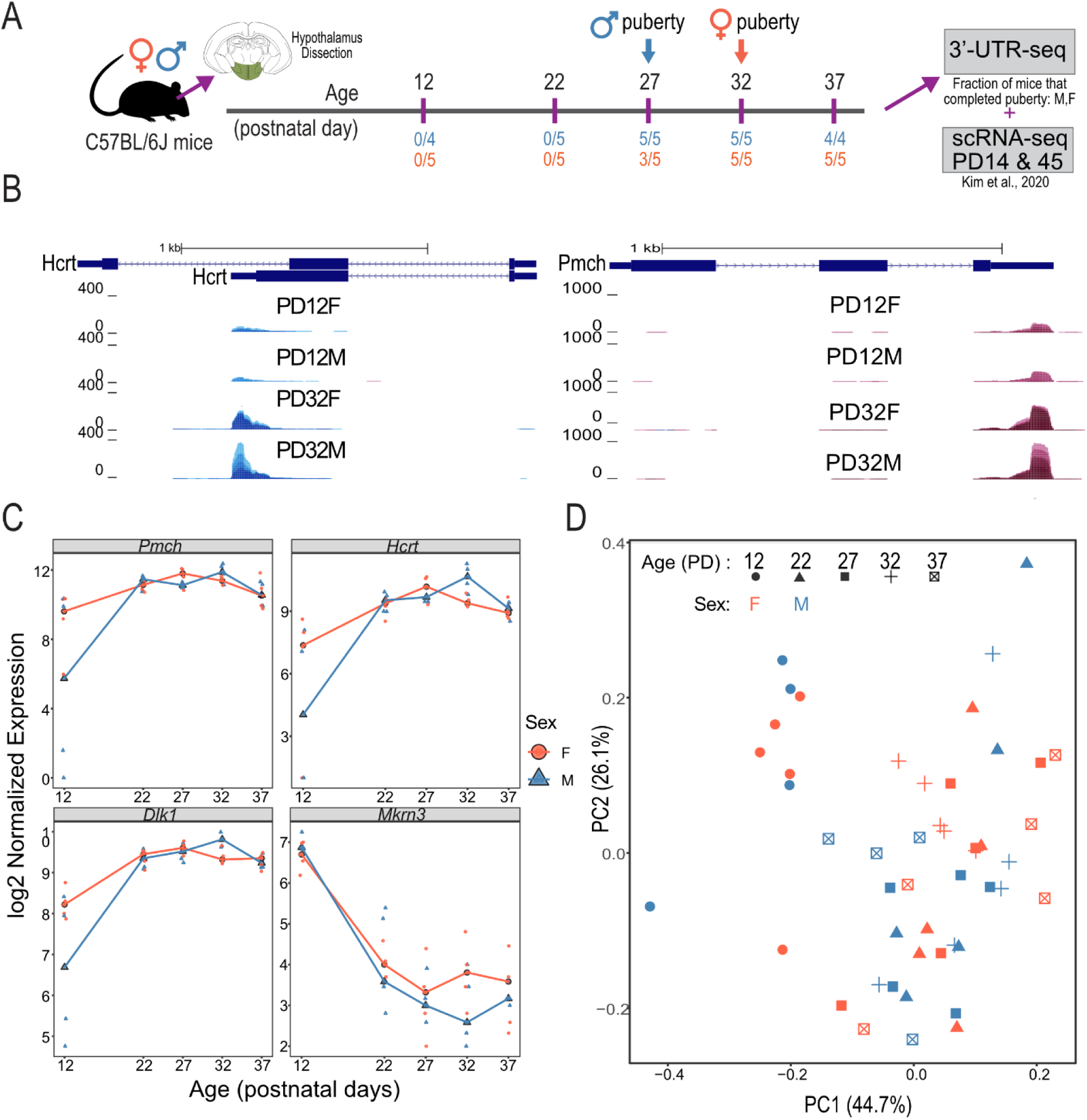
Overview of the hypothalamic mouse transcriptome at five timepoints across pubertal development in males and females. A) Schematic of samples taken across pubertal development. Whole mice hypothalami were dissected at postnatal days (PD) 12, 22, 27, 32, and 37 in both male and female C57BL6/J mice. Arrows dictate the average age of puberty in males and females, respectively. Extracted hypothalamus samples underwent 3’UTR RNA-seq. B) Genome browser of the *Hcrt* (top) and *Pmch* (bottom) 3’UTR at PD12 and PD22 in males and females. C) Distribution of normalized counts of *Pmch, Hcrt, Dlk1,* and *Mkr3* at every age and timepoint. The X-axis is age, and the Y-axis is log2-transformed RUVseq and ERCC-spike in normalized counts. Red lines and circles represent female samples, while blue lines and triangles represent male samples. D) Principal component analysis (PCA) of normalized gene expression across all samples and ages. The first two PCs are plotted with sexes designated with colour and ages designated by shape.

We first investigated the gene expression of four genes whose gene expression and expression patterns are well characterized in the hypothalamus, namely *Mkrn3, Cartpt, Dlk1,* and *Pomc*, and verified that our analyses captured these previously reported expression dynamics in the hypothalamus [13, 70–73] (Figure 1B, C). We next leveraged 183 genes where we completed qPCR in the same mice [10] as in our RNA-seq data to correlate the gene expression of every gene in each sample between the two technologies. Samples were highly correlated between the RNA-seq and qPCR data based on these 183 overlapping genes (R^2^ mean = 0.698, sd = 0.0270) (Supplementary Figure S1A). As previously shown with qPCR of selected puberty-related genes, PCA of the RNA-seq data revealed the greatest overall change in gene expression between PD12 and all other timepoints in both male and female hypothalamus samples (Figure 1C). Lastly, we demonstrated that every sample in our 3’UTR-seq data is highly correlated to one another, with Pearson’s correlation coefficient ranging from 0.89 to 0.99 between samples (0.98-0.99 between replicates) (Supplementary Figure S1B).

### Pairwise differential gene expression across pubertal development reflects the hypothalamic cellular composition dynamics and puberty-relevant transcriptional control

To investigate the transcriptomic dynamics in the hypothalamus throughout pubertal development, we identified DEGs between the studied age groups in male and female mice separately, as well as DEGs between sexes at each timepoint (FDR adjusted p-value < 0.05 and absolute fold-change > 1.5). We denoted age-biased DEGs with a positive fold-change to have higher expression in the later timepoint in development (e.g., PD12 vs. PD22, PD22 is greater). When comparing sexes, we denoted DEGs with a positive fold-change to have higher expression in females than males. Our RNA-seq data and results of differential analysis can be visualized and downloaded interactively using our Shiny App (https://wilsonlab-sickkids-uoft.shinyapps.io/hypothalamus_gene_shiny/).

We found that most DEGs are established before the physical signs of pubertal onset (i.e., vaginal opening in females, between separation in males) between PD12 and PD22, with 32% (511/1560) of DEGs overlapping between sexes (Figure 2A), with the fold-changes of all 1560 genes being highly correlated (R^2^ =0.905). Accordingly, we considered the transcriptomic differences between PD12 and PD22 to be conserved across sexes. We interrogated these DEGs using ActivePathways, a method that integrates the p-values of DEGs in males and females to identify enriched pathways in a sex-aware manner [43]. Upregulated DEGs enriched for pathways related to glial-cell development, particularly myelination in males (Figure 2B). While the transition from oligodendrocyte precursor cells (OPCs) into mature oligodendrocytes (MOs) has been characterized in post-pubertal mice [74], oligodendrocyte development before and during puberty is not well characterized. Downregulated DEGs tended to enrich pathways involved in cell differentiation, cell morphogenesis, and proliferation (Figure 2B), likely reflecting how the brain, including the hypothalamus, doubles in volume between PD12 and PD20 [75].

**Figure 2.**
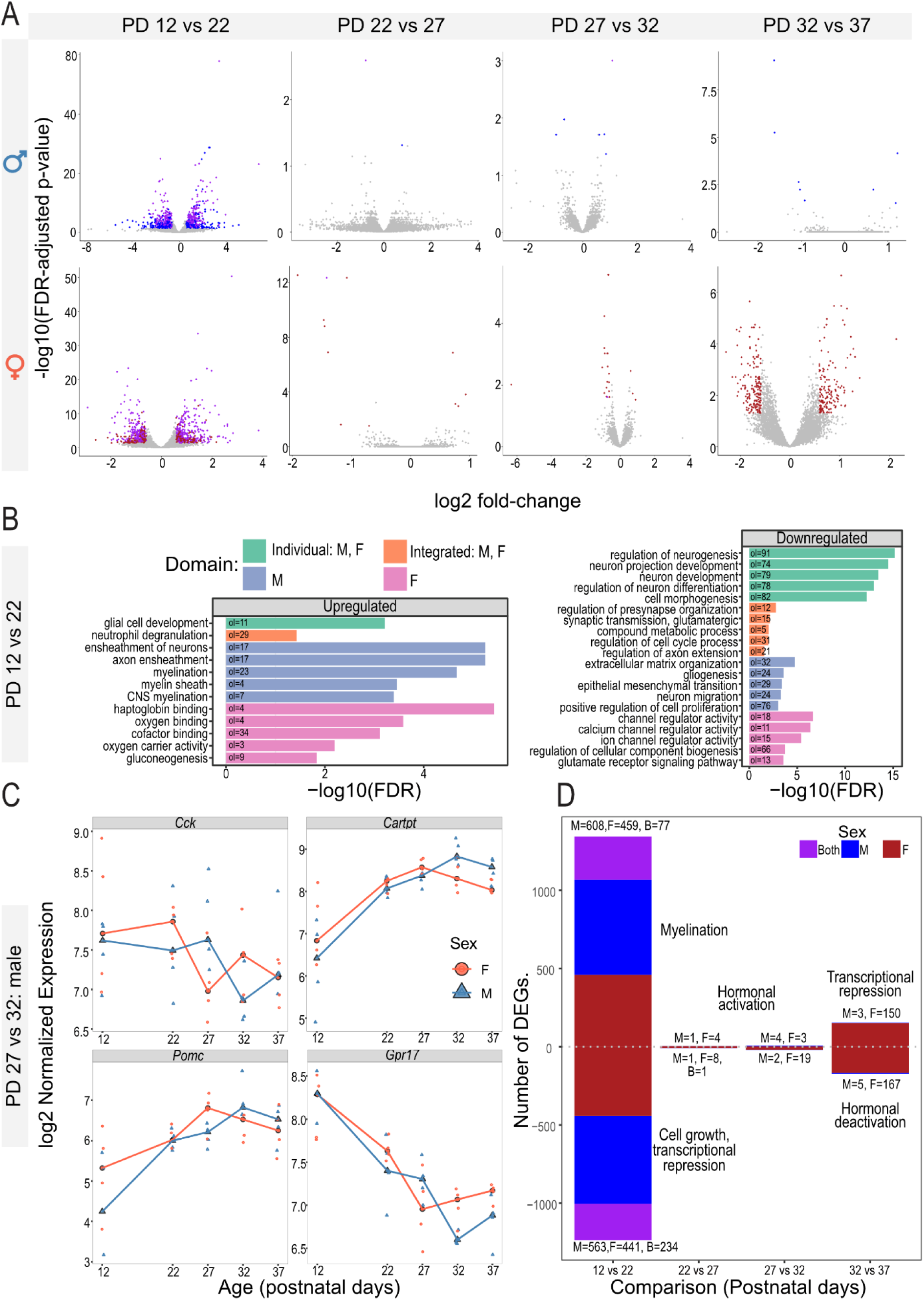
Differentially expressed genes (DEGs) across postnatal development in the mouse hypothalamus. A) Volcano plot of differentially expressed genes in each pairwise timepoint. The X-axis is the log2(fold-change) of the DEG, and the Y-axis is the -log10 (FDR-adjusted P-value) of the DEG as identified by DESeq2. Genes in grey are not detected as DE (FDR-adjusted P-value < 0.05, absolute fold-change > 1.5). Genes in blue are DE in males, genes in red are DE in females, and genes in purple are DE at the same timepoint in both males and females. B) Barplot of enriched pathways derived from DEGs between PD12 and PD22 in male and female mice. Genes are separated by upregulated and downregulated DEGs. Barplots show the - log10(FDR-adjusted P-value) of enrichment. Green bars represent pathways detected in both sexes, orange bars represent pathways detected by integrating sexes, blue bars represent male-driven pathways, and pink bars represent female-driven pathways. C) Expression profile of the four remaining DEGs. D) Barplot summarizing the number and major theme of pairwise DEGs across each timepoint. The X-axis is each timepoint, and the Y-axis is the number of DEGs. Positive genes were older-biased, and negative genes were younger-biased. D) Expression profile of DEGs involved in hypogonadotropic hypogonadism. The X-axis is age, and the Y-axis is log2-transformed RUVseq and ERCC-spike in normalized counts. Red lines and circles represent female samples, while blue lines and triangles represent male samples. Expression profile of differentially expressed GWAS genes (right). Row-normalized heatmap of GWAS-associated genes that were also detected as differentially expressed in our RNA-seq data Rows are genes, and columns are samples.

We detected 317 DEGs in post-pubertal female timepoints, however this transcriptomic signature was not found in males (PD32 vs. PD37) (Figure 2A). We identified several downregulated puberty-relevant neuropeptides, including *Tacr1* [76] and *Sst* [77]. Upregulated DEGs included genes encoding transcriptional regulators that regulate genes involved in the hypothalamus-pituitary-gonadal axis (e.g., *GnRh*, *Lhb*, *Ar*, and *Pgr*) such as *Cited2* [78], *Fgfr2* [79], *Lcor* [80], and *Sp1* [81] (Supplementary Figure S2A, B). We reasoned that the increase in the expression of transcriptional regulators paired with a decrease in neuropeptide expression after puberty might reflect the well-documented release and return of transcriptional repression upon activation of pubertal development [12]. Interestingly, female DEGs that decrease in expression before vaginal opening (PD12 vs. PD22) are overrepresented in female DEGs that increase after puberty (PD32 vs. PD37; 23 genes, FDR-adjusted p-value = 5.06 x 10^-12^). Pathway analysis of these overlapping genes enriched for “transcriptional co-repression activity” (*Skil*, *Wwtr1*, *Cited2*) (3 genes, FDR-adjusted p-value = 0.044) and pathways involved in cell and tissue development (Supplementary Figure S2C, Supplementary Figure S3). Consistent with the hypothesis of female release of hormonal repression, females contained 8 upregulated DEGs before puberty (PD12 vs. PD22) mapping to the “response to peptide hormone” gene ontology (8 genes, FDR-adjusted p-value < 0.01), including *Th*, which expresses thyroid hormone and regulates the hypothalamus-pituitary-adrenal axis, and *Agrp*, which modulates puberty with leptin (Supplementary Figure S3) [82].

We identified a relatively small group of DEGs spanning the ages when physical signs of puberty emerge (PD22 vs PD27: number of DEGs in males (N_male_) = 2, number of DEGs in females (N_female_)=12 and PD27 vs PD32: N_male_ = 6, N_female_ = 22). Strikingly, 5 of the 6 DEGs found in males between PD27 and PD32 were puberty-relevant neuropeptides, including downregulated Cholecystokinin (*Cck*) [83], and upregulated CART peptidase (*Cartpt*) [71], Pro-melanin concentrating hormone (*Pmch*) [84], Orexin (*Hcrt*) [85, 86], and Proopiomelanocortin (*Pomc*) genes [73] were upregulated (Figure 1C, Figure 2C, D). *Hcrt* [85, 86], Oxytocin (*Oxt*) [87], and *Axl* [88], whose knockout leads to pubertal delay in mice, are all differentially expressed during puberty in female mice and peak in expression at PD27 upon vaginal opening (PD22 < PD27, PD27 > PD32; Figure 2). Accordingly, the relatively few DEGs in this age window (PD22 vs PD27, PD27 vs PD32) consist of puberty-relevant neuropeptides that peak in expression during the average age of pubertal onset in both males and females.

Lastly, unlike the PD12 to PD22 age window, we found relatively few sex differences at any timepoint (N_male_=41, N_female_=22), with almost half of all sex-biased DEGs at any timepoint mapping to a sex chromosome (chrX=5, chrY=21) (Supplementary Figure S4A,B). While there are fewer sex differences than DEGs across timepoints, four puberty-relevant genes are sex-biased. Specifically, *Tcf7l2* [89] is female-biased at PD27, and *Etnppl*, *Cryab* [20], and *Hcrt* [85, 86] are male-biased at PD32 (Supplementary Figure S4C).

### Age-biased differentially expressed genes compared to human RNA-seq identifies conserved modules of development and post-natal cellular growth

We then tested whether the one-to-one orthologs these up- and down-regulated DEGs between PD12 and PD22 related to comparable DEGs in human RNA-seq datasets using the Differential Expression Enrichment Tool (DEET) [46]. Briefly, DEET stores the DEGs from 3162 uniformly processed and analyzed comparisons from The Cancer Genome Atlas (TCGA), Genotype-Tissue Expression Consortium (GTEx), and 142 studies within SRA, including hundreds of brain-related comparisons [46–48, 90].

Upregulated DEGs between PD12 and PD22 enriched for comparisons related to cellular composition, glioma development, and comparisons investigating biological processes regulated by the hypothalamus (e.g., Body-mass-index, body temperature, blood pressure) in the pituitary gland in GTEx (Supplementary Figure S5A) [47, 48]. In accordance with the traditional pathway enrichment, we found that many of the genes driving the enrichment of the GBM and pituitary studies are related to myelination (Supplementary Figure S5B-D). Similarly, gene expression differences between oligodendrogliomas vs. astrocytomas [91] were strongly correlated DEGs (up- and down-regulated) to our PD12 vs. PD22 comparison in males (R^2^ = 0.400, FDR = 9.07 x 10^-16^) (Supplementary Figure S6), consistent with our observation of the activation of oligodendrocyte-growth genes. Downregulated DEGs show strong enrichment of studies related to cellular growth and differentiation and drug treatments of neuronal stem cell lines, which may also be related to the substantial growth of the mouse brain between PD12 and PD22 (Supplementary Figure S5E).

DEET also identified that the PD12 vs. PD22 age-biased DEGs in our mice were significantly associated with DEGs comparing infant and child males in the pre-frontal cortex (PFC), suggesting that our study in mice captured a set of genes related more generally to mammalian postnatal brain development (119 genes, FDR = 8.87 x 10^-18^, R^2^ = 0.723, FDR = 1.42 x 10^-17^) [92] (Figure 2E-G, Supplementary Figure S6B). Enriched biological pathways for genes overlapping with infant vs. child in the PFC were primarily related to neuron differentiation (Supplementary Figure S6B). Additionally, we found that transcription factors and chromatin regulators (e.g., *DNMT3A*, *YBX1*, *SOX11*, *SOX4*, *TOP2A,* and *TOP2B*) were younger-biased in both species, while *MAL* had the strongest shared increase in expression in both mice and humans. The conserved increase in *MAL* may also reflect the sex-biased increase in white matter found in humans during postnatal development and adolescence [93]. In contrast, the 30 shared DEGs between Adolescent vs. adult males in the PFC were also highly associated to our PD12 vs. PD22 comparison (R^2^ = 0.878, FDR = 1.38 x 10^-16^) [92] (Supplementary Figure S6C) were involved in calmodulin binding and chemical synaptic transmission (Supplementary Figure S6C). Together, these results show that a substantial proportion (140/1171) of the genes involved in early postnatal developmental programming in the mouse hypothalamus is conserved across species and tissue.

### Human puberty genome-wide association study candidate genes are differentially expressed before and after puberty in the hypothalamus

We investigated if the DEGs we measured before and after pubertal development overlapped with a set of candidate human genes associated with the age of menarche in females and voice breaking in males (i.e., pubertal timing) from GWAS analysis of the U.K. Biobank [1, 5, 10] to further link our transcriptional dynamics to puberty in humans (Supplementary Figure S7). Interestingly, we found that DEGs detected before and after the physical onsets of puberty in males and females and after the onset of puberty in females, but not during puberty in either sex were overrepresented within these GWAS genes (PD37-female > PD32-female: FDR = 0.00520, and odds-ratio = 2.88, PD12-female > PD22-female: FDR = 0.0423 and odds ratio 1.72, PD12-male > PD22-male: FDR = 0.0685 and odds ratio 1.54) (Supplementary Figure S7). Moving from GWAS genes, rare-disease genes that lead to precocious or delayed puberty, we found that three genes implicated in hypogonadotropic hypogonadism (HH) [4] were differentially expressed between PD12-PD22. Specifically, *Il17rd* is downregulated in males and females (PD12 vs. PD22), *Sema3e* is downregulated in females but not males, and *Rab3gap1* is upregulated in males and females (PD12 vs. PD22) (Supplementary Figure S7). Briefly, *Il17rd* is a member of the interleukin-17 receptor protein family and is important in regulating growth through fibroblast growth factor and MAPK/ERK signaling [94]. *Sema3e* is a semaphorin, which acts as axon guidance ligands and organogenesis [95]. *Rab3gap1* is a member of the Rab3 protein family where it’s involved in endoplasmic reticulum structure and has also been implicated in the proper development and migration of neurons [96]. *Il17rd*, *Sema3e*, and *Rab3gap1* were not differentially expressed at other timepoints. Together, our overrepresentation of pre- and post-pubertal DEGs with human puberty-GWAS genes, as well as our overlap of pre-pubertal DEGs with HH disease genes, suggests that the DEGs detected before and after puberty may be indirectly involved in pubertal regulation in both mice and humans.

### Genes expressing metabolic and reproductive neuropeptides display an age-by-sex interaction in gene expression along the pubertal transition

To identify genes whose expression is conditional on both age and sex, we leveraged the varimax rotated principal component analysis (vrPCA) (Figure 3A, See Materials and Methods for details). We were particularly interested in vrPCs associated with an age-by-sex interaction because of the known sex bias in pubertal onset and development in both mice and humans. As such, puberty-relevant genes and pathways would have slightly offset or divergent age-biased gene expression patterns. Accordingly, four vrPCs were associated with age, one with sex, and one with an age-by-sex interaction (Figure 3A). The scores of the vrPC conditional on age and sex, vrPC 16, were dynamic between PD12 and PD27 and showed sex bias at PD32 (Figure 3B, C). For simplicity, we denote the genes associated with vrPC16 as age-by-sex associated genes.

**Figure 3.**
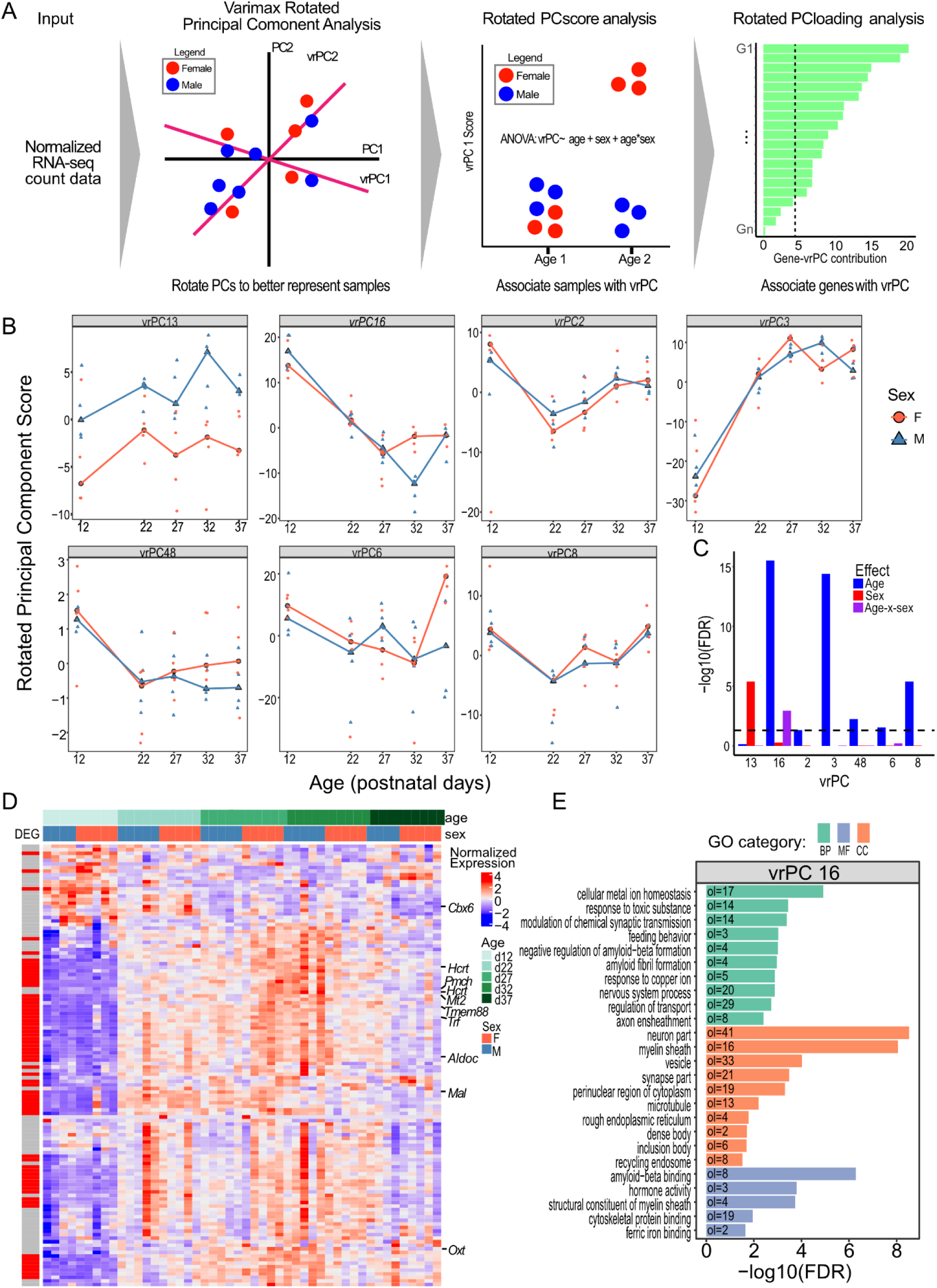
Evaluation of varimax-rotated principal component analysis revealed genes involved in sex-by-age interactions. A) Schematic of how varimax rotated PCA is applied to our data. B) Distribution of scores for enriched vrPCs. Male samples are triangles and blue lines, and female samples are circles and red lines. vrPC16 is highlighted because the genes associated with this vrPC are focused on for the rest of this study. C) Barplot showing the association between each significant varimax rotated PC (vrPC), age, and sex. The X-axis shows vrPCs whose scores are associated with age, sex, or an age-by-sex interaction (7/48 total vrPCs). Red bars show the significance of sex, blue bars show the significance of age, and purple bars show the significance of an age-by-sex interaction. D) Heatmap of the gene expression patterns of genes associated with vrPC16. Each row is a gene, and each column is a sample. The heatmap is populated by the log2-RUV-seq normalized gene expression of each gene. Rows are annotated by whether the gene displays pairwise expression in at least one pairwise timepoint. Columns are annotated by age and sex. E) Barplot of enriched pathways derived from genes strongly associated with rotated PC 16. Barplots show the -log10(FDR-adjusted P-value) of enrichment. Green bars represent pathways deriving from gene-ontology biology pathways, red bars represent pathways deriving from gene-ontology cellular components, and blue bars represent pathways deriving from gene-ontology molecular functions.

In total, we identified 129 age-by-sex associated genes, 66 of which were differentially expressed between PD12 and PD22 in males or females (Figure 3). Interestingly, the four genes with the strongest association with an age-by-sex interaction based on their vrPC loading are all hormone-producing genes that have been linked to pubertal regulation or dynamics: *Pmch*, *Hcrt*, *Oxt,* and *Trf* [84, 85, 97, 98] (Figure 3D). While these genes with top loadings shared similar expression patterns (i.e., a secular increase in gene expression from PD12-PD27 before diverging by sex), 21 genes, including puberty-regulating *Cbx6* [12], a member of the Polycomb repressive complex, decrease in gene expression before diverging by sex (Figure 3D). Pathway enrichment of age-by-sex associated genes were enriched for hormone activity (precision = 0.100, FDR = 1.60 x 10^-4^), negative regulation and transmission of nerve impulse (precision = 0.167, FDR = 0.0248), and neuron and oligodendrocyte development pathways, including “neuron part” (precision = 0.339, FDR = 2.99 x 10^-9^) and “myelin sheath” (precision = 0.132, FDR = 8.82 x 10^-8^) (Figure 3E).

Likewise, DEET analysis of age-by-sex associated genes most strongly enriched for human DEG comparisons influencing glial cell growth, namely comparing glioblastoma subtypes and LGG drug treatments in the TCGA database, and neuronal-controlled disorders in relevant tissues, namely sporadic amyotrophic lateral sclerosis in motor neurons and individuals with schizophrenia in the adrenal glands from the GTEx database (Supplementary Figure S8A). Interestingly, unlike in PD12 vs. PD22 comparisons, these age-by-sex associated genes also enriched for many relevant comparisons in the hypothalamus, including age and body mass index (BMI) from the GTEx database (Supplementary Figure S8B). The genes driving the enrichment of these hypothalamus comparisons were predominantly puberty-relevant hormonal neuropeptides with a high vrPC16 loading, namely *OXT, AVP, HCRT,* and *PMCH* (Supplementary Figure S8C,D).

Recently, spatially resolved single-cell transcriptomics have been performed along the pubertal transition of the female rat arcuate nucleus [16]. They identified three gene-expression modules associated with the pubertal transition. Broadly, they categorized genes associated with these modules as: module 1) glial cell enhancement and neuron proliferation in response to estradiol, module 2) hormone secretion, and module 3) neuronal differentiation and signal transmission [16]. The age-by-sex associated genes we identified were over-represented in all three modules (module 1: p-value = 2.23 x 10^-13^, odds-ratio = 6.80, genes = 29; module 2: p-value = 0.0506, odds-ratio = 1.97, genes = 11; module 3: p = 4.11 x 10^-11^, odds-ratio = 4.22, genes = 38). Together, genes associated with an age-by-sex interaction across puberty are involved in hypothalamic hormonal activity, neuronal development, and oligodendrocyte development.

### Cellular composition of the postnatal hypothalamus

The hypothalamus exhibits considerable cellular heterogeneity reflecting its multimodal functions [21, 23]. To characterize the cell type-specific underpinnings of pubertal development in the hypothalamus, we integrated scRNA-seq in the hypothalamus with our temporal bulk RNA-seq. We leveraged data from Kim et al., 2020, which contained scRNA-seq from the mouse hypothalamus before and after puberty (PD14 and PD45) [23]. We incorporated the cell-type labels provided by Kim et al., 2020 (hypothalamic neurons, oligodendrocytes, tanycytes, ependymal cells, astrocytes, microglia, and endothelial cells) [23] with cell-type identification analysis of clusters measured with Seurat [53] (see Materials and Methods for Details). Our cluster analysis further subdivided oligodendrocytes into OPCs, DOs, and MOs. It also subdivided neurons into neurons and neuroendocrine cells (Figure 4).

**Figure 4.**
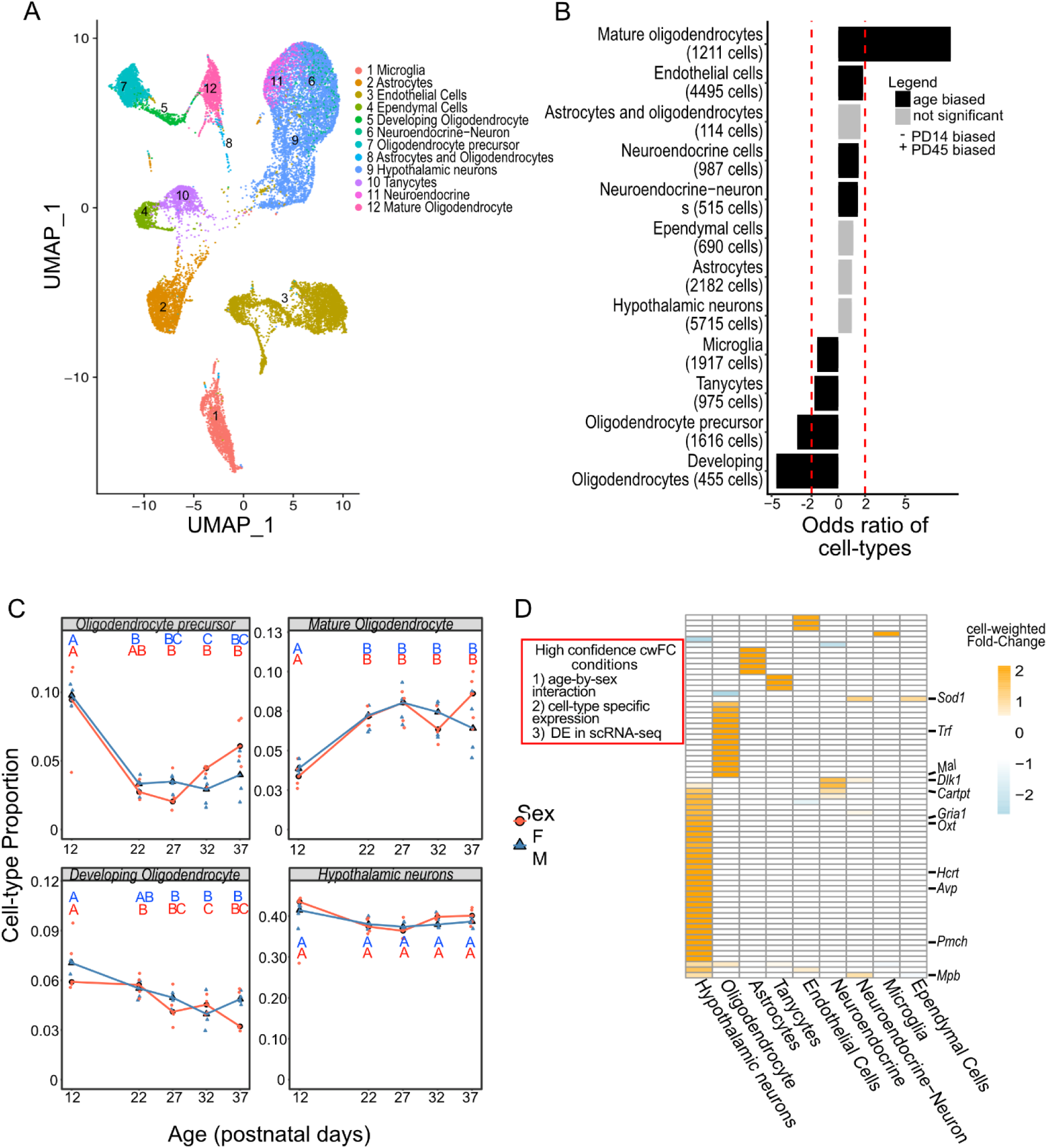
Cell-type specific gene expression across the developing hypothalamus. A) Lower-dimension representation of scRNA-seq data in the PD14 and PD45 mouse hypothalamus with the Uniform Manifold Approximation and Projection (UMAP). Cell labels were identified using a mixture of labels provided by Kim *et al.,* 2020 and unsupervised clustering. B) Distribution of cell proportions estimated from RNA-seq deconvolution at each age and time-point. The X-axis is age, and the Y-axis is estimated cell-type proportions. Red lines and circles represent female samples, while blue lines and triangles represent male samples. Letters represent significance using a Tukey post hoc test after identifying differences in cell-type proportion with ANOVA. C) Barplot of cell-type proportion differences within each cluster (Fisher’s exact-test). Red bars designate a fold-change of two between ages. Each column is a cell-type with the number of DEGs mapping to that cell-type in brackets. D) Heatmap of gene-normalized cell-weighted fold-changes (cwFold-changes) of the 129 age-by-sex associated genes and are DE in the complementary direction in the scRNA-seq data.

We first investigated hypothalamic cell-type proportion dynamics across pubertal timepoints. When investigating the scRNA-seq data alone, we found that oligodendrocytes were the most dynamic cell types across puberty (Figure 4B), with MOs increasing in proportion over time (PD14 < PD45) (Bonferroni-adjusted p-value = 1.90 x 10^-106^, fold-change = 8.43), and OPCs (Bonferroni-adjusted p-value = 3.94 x 10^-99^, fold-change = -3.07) and DOs (Bonferroni adjusted p-value = 8.18 x 10^-56^, fold-change = -4.65) decreasing in proportion over time (PD14 > PD45). There was also a lesser but significant increase in endothelial (Bonferroni-adjusted p-value = 1.09 x 10^-54^, fold-change = 1.856) and neuroendocrine cell (Bonferroni adjusted p-value = 4.81 x 10^-7^, fold-change = 1.52) proportions over time.

Next, we used estimated hypothalamic cell-type proportions in our bulk RNA-seq data and RNA-seq deconvolution, mapping cell-type proportion changes across our developmental trajectory. Benchmarking RNA-seq deconvolution in the hypothalamus is important because it has both highly similar cell types (e.g., neuron vs. neuroendocrine) and highly distinct cell types (neuron vs. endothelial cell) amongst its many total cell types. To find the most reliable RNA-seq deconvolution tool in our system, we compared the cell-type proportions of nine different RNA-seq deconvolution tools [25, 26, 60–65] to the scRNA-seq data (See Materials and Methods for Details), where we found that the NNLS-MuSiC tool was the most accurate method, a method that has previously performed well on brain tissue [99] (Supplementary Table S2). As in the scRNA-seq data, we found that MOs increase in cell-type proportion until puberty, and OPCs (p-value = 3.68 x 10^-11^) and DOs decrease in cell-type proportion until puberty (Figure 4C, Supplementary Figure S9). These results show that oligodendrocytes expand from OPCs into MOs during puberty.

### Age-by-sex associated transcriptional dynamics map to genes involved in neuropeptide activation and oligodendrocyte maturation.

We next used these scRNA-seq data to assign age-by-sex associated genes to their cell type of origin using scMappR [24], using both the PD14 and PD45 timepoints in the scRNA-seq data [23]. Overlapping the 129 cell type-specific age-by-sex associated genes with cell type-specific DEGs from the scRNA-seq data [59] (PD14 vs. PD45) yielded a set of high-confidence cell-type specific genes (n=67), whose gene expression patterns are conditional on both age and sex (Figure 4D). Four high-confidence genes with the top age-by-sex loadings mapped to neurons, namely *Pmch*, *Hcrt*, and *Oxt,* while *Trf* mapped to oligodendrocytes [74] where it may act as a cofactor for iron in myelination [100]. *Dlk1* and *Gria1*, which have previously been implicated in pubertal disease and ovulation rate respectively [101, 102], mapped to neuroendocrine cells and neurons (Figure 4D, Supplementary Figure S10). Next, we investigated whether the neuron- and neuroendocrine-mapping age-by-sex associated genes overlapped with translated mRNA in lepRb+ neurons in the hypothalamus [51] using Trap-Seq because leptin is a functional activator of pubertal initiation [20, 71, 82, 103, 104]. Interestingly, 21 of our neuron- and -neuroendocrine-mapping genes were enriched in lepRb+ neurons (p-value = 4.33 x 10^-9^, odds ratio = 5.96) (Supplementary Figure S11). These genes included *Cartpt*, *Dlk1*, and *Sod1*, which can all influence pubertal timing or fertility [71, 101, 105, 106], showing that many of our neuronal-mapping genes containing puberty-relevant transcriptional dynamics are expressed in cell-types that are responsive to known pubertal activators. Overall, genes with transcriptional dynamics conditional on age and sex were cell type-specific, puberty-relevant, and often related to neuron and oligodendrocyte development.

### Lineage reconstruction of oligodendrocytes links pubertal genes to cellular development

We observed an active transition of oligodendrocytes from OPCs into MOs in the postnatal hypothalamus (Figure 2B, D, Figure 4), and detected clear manifold from OPCs to DOs and MOs in the reprocessed scRNA-seq data (Figure 4A). We next set to investigate transcription factors (TFs), age-by-sex associated genes, and puberty-GWAS genes that map to oligodendrocyte expansion. By using Slingshot, a bioinformatic package that identifies cellular lineages across cell types [67], we measured a pseudotime trajectory from OPCs to MOs (Figure 5A). We then applied the tradeSeq [68] pipeline and the “Association Test” to identify genes that are associated with this trajectory (FDR < 0.05, fold-change > 1.5).

**Figure 5.**
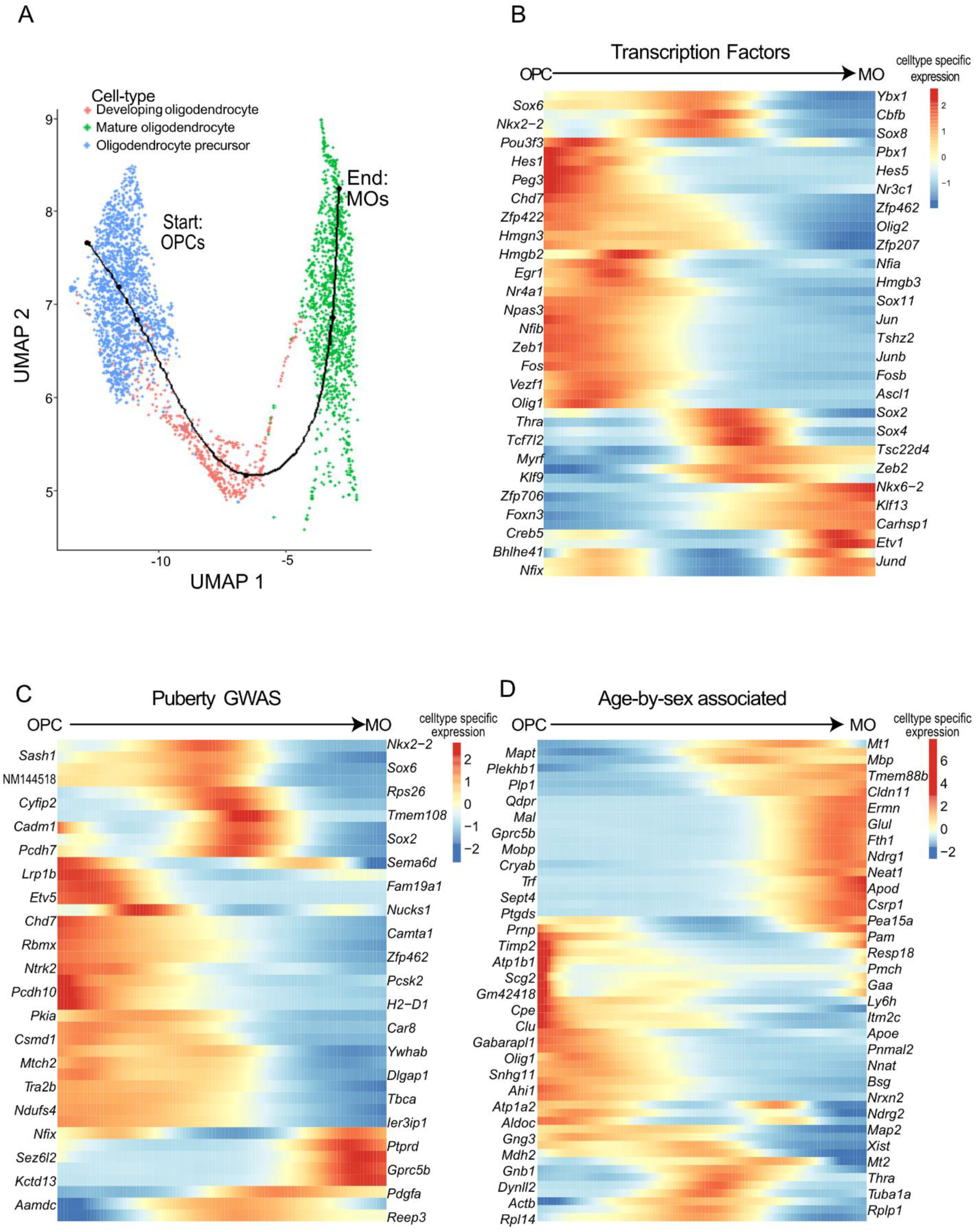
Pseudotime of hypothalamic oligodendrocyte development. A) Lower-dimension representation of oligodendrocyte scRNA-seq data in the PD14 and PD45 mouse hypothalamus with the Uniform Manifold Approximation and Projection (UMAP) overlaid with the pseudotime trajectory identified with Slingshot. Points are cells coloured by cell-type. and the line is the plotted pseudotime trajectory measured with Slingshot, starting with OPCs and plotted with the tradeSeq R package. B) Heatmap of transcription factors associated with pseudotime. C) Heatmap of puberty-associated GWAS genes associated with pseudotime. D) Heatmap of oligodendrocyte-mapped age-by-sex associated genes associated with pseudotime. For a gene to be included, it must be associated with an age-by-sex interaction (i.e., varimax 16), mapping to oligodendrocyte precursor cells, developing oligodendrocytes, or mature oligodendrocytes with scMappR, and associate with pseudotime. For B-D, rows are genes associated with pseudotime. Columns are portions of the pseudotime trajectory blocked into 200 smoothers using tradeSeq. Heat is measured by scaling the predicted smoothers with the scale function in R.

Overall, we found 1294 genes to be associated with pseudotime and found that these oligodendrocyte-pseudotime genes overrepresented puberty-GWAS genes (FDR = 4.6 x 10^-3^, 27 genes). Oligodendrocyte-pseudotime TFs and puberty-GWAS genes whose expression peaks in in DOs include genes involved in thyroid hormone response (*Nkx2-1* and *Thra*) and cell differentiation and development (*Sox2*, *Tcf7l2*, *Egr1*, *Hes1*) (Figure 5B,C). Interestingly, thyroid hormone, which can impact pubertal timing and function [14], also plays a direct role in OPCs expanding into MOs [107], which may explain the transcriptional activity of *Thra* in DOs. Most TFs and candidate GWAS-associated genes peaked in expression in OPCs. These genes, while potentially involved in puberty, may be involved in the differentiation of many cell types including those in the hypothalamus. For example, *Sox2* peaks in expression at OPCs and functions as a key marker for cellular differentiation in general [108, 109], and mutations in *Sox2* lead to developmental abnormalities throughout the entire body, including HH [110–112]. Accordingly, we hypothesize that many of these TFs and puberty-GWAS genes would be expressed during the differentiation of many cell types, not just in hypothalamic oligodendrocyte expansion.

We next overlapped the age-by-sex associated genes that mapped to oligodendrocytes with the 1294 oligodendrocyte-pseudotime genes and found that the gene lists significantly overlapped (FDR = 4.6 x 10^-3^, 58 genes). Of the 58 overlapping genes, we found an even distribution of genes mapping to OPCs, DOs, and MOs (Figure 5D). These genes included core oligodendrocyte stage-specific regulated proteins such as *Mbp*, *Mobp*, *Mal*, and *Olig1* [113], puberty and HPA-linked *Thra*, melatonin receptors *Mt1* and *Mt2*, and *Pmch (*Figure 5D). Three of these genes, namely *Sox2*, *Chd7*, and *Stub1*, can lead to HH in humans [4]*. Chd7* works with *Sox10* to promote myelination by co-occupying and promoting the expression of myelinogenic genes [114], and *Stub1* has a less studied role in oligodendrocytes. Two transcriptional regulators whose mutations lead to both HH and hypomyelination are *Polr3a* and *Polr3b* [115]. We found that other members of the polymerase 3 complex, namely *Polr3e* and *Polr3h*, were identified as oligodendrocyte-pseudotime genes (Supplementary Figure S12). Together, oligodendrocyte-pseudotime genes previously detected as puberty-GWAS genes tend to be involved in the differentiation of many cell types, while oligodendrocyte-pseudotime genes overlapping with age-by-sex associated genes suggest a more varied relationship between oligodendrocyte expansion and puberty.

## Discussion

The hypothalamus is responsible for pubertal initiation and is a component of the network that modulates pubertal timing. The regulation of puberty (onset and progression) is a dynamic, non-linear process, making it imperative to examine multiple timepoints before, during, and after puberty. Puberty is also inherently sex-biased in initiation, regulation, and manifestation, resulting in sex also being an important variable to include when studying puberty. Combining our bulk RNA-seq spanning of 5 timepoints in male and female mice with publicly available data from GWAS of pubertal timing; lists of genes that, when mutated, cause absent, delayed or precocious puberty; and hypothalamic scRNA-seq of pre-(PD14) and post-pubertal (PD45) mice [23] enabled us to replicate previous findings and identify new genes and cell-types involved in hypothalamic regulation of puberty.

### Transcriptome-wide gene expression dynamics in the hypothalamus reflect hormonal activation and epigenetic control

Our neuron- and neuroendocrine-mapping transcriptomic dynamics reflected previously reported mechanisms of pubertal regulation while implicating new genes involved in puberty. Differential gene expression in male and female mice during puberty (i.e., PD22-PD27, PD27-PD32) primarily displayed differences in genes coding for pubertal and metabolic hormones and neuropeptides, namely *Oxt*, *Hcrt*, *Cartpt*, *Avp*, *Cck*, *Pmch*, and *Pomc* [71, 73, 83–85, 87, 116], with all these genes other than *Cck* and *Pomc* also displaying an age-by-sex interaction. Transcriptional regulators in females decrease in expression before puberty and increase in expression afterward while peptide hormone-producing genes such as *Hcrt, Oxt*, and *Th,* peak in expression at PD27 in females. In males, the decrease in transcriptional repression is less pronounced and these genes peak in expression at PD32.

Inactivating mutations in *Hcrt* leads to narcolepsy type-one, which has a high association with precocious puberty in both male and female humans [86, 117]. Orexin, produced by *Hcrt*, has been shown to have an inhibitory effect on Gnrh in ovariectomized female mice [118]. In male rats, orexin injections are associated with a decrease in *Gnrh*, *Kiss1*, *NPy* expression paired with a decrease in reproductive behaviors [119]. Despite these findings, the mechanistic and potentially sex-specific role of orexin on pubertal inhibition remains unclear [120]. *Oxt* encodes oxytocin and plays an important role in parturition, lactation, social behavior, and parental behavior. Oxytocin may influence pubertal activation in a number of ways, such as influencing the estrogen receptor in female mice [121], interacting with Prostaglandin E2 in astrocytes to stimulate GnRH neuron in female mice [87], and through direct stimulation of spermatogenesis in the mouse testes. This mechanism is also bi-directional, as oxytocin levels during puberty influence adult behavior in a sex-specific manner [121].

The inverse relationship between epigenetic repressors and hormonal activation during puberty in females is consistent with the epigenetic regulation of hypothalamic hormones during puberty, which is known to be driven by the release of transcriptional repression in female rats [9, 12]. Interestingly, we did not find this pattern in males, however given the peak of *Oxt, Hcrt*, and *Th* were found at PD32, perhaps these genes would have increased in expression at a later timepoint in males (e.g., PD42). Lastly, we found that neuron- and neuroendocrine-mapping genes associated with an age-by-sex interaction tended to be previously implicated in puberty (Figure 4, Figure 5D). These findings provide further support to previously implicated pubertal regulators and suggest that lesser-known candidate genes identified using the same approaches are involved in same regulatory network [122].

Three puberty-relevant, neuronally-mapped age-by-sex associated genes highlighted in this study that have been not as thoroughly studied as the neuropeptide producing genes mentioned above were *Gria1*, *Dlk1*, and *Cartpt*. *Gria1* is a primary receptor of glutamate, a neurotransmitter that directly messages GnRH neurons [2, 102]. Recent studies in female rats show that *Gria1*, along with a network of genes implicated in the epigenetic control of puberty, is under the shared regulation of *Kdm6b* at puberty in the hypothalamus [11]. Genetic variants in *Dlk1* are associated with CPP and *Dlk1* is highly associated with age of menarche through GWAS [1, 72, 105, 123]. While not evaluated experimentally, *Dlk1*’s ability to regulate Notch signaling has been a proposed mechanism to control pubertal timing. Our lab previously completed *Dlk1* expression dynamics with the same RNA using qPCR [10] in multiple different tissues, finding the same pattern of age-biased gene expression between PD12 vs. PD22 expression in the male and female, sex bias across puberty in the pituitary gland, and a decrease in expression over time in the testis and ovaries [10], further suggesting the cell- and tissue-specific roles of *Dlk1* in pubertal regulation. Lastly, *Cartpt* plays a core role in the function of CART neurons, a neuronal subtype that receives signals from leptin and alters pubertal timing in female mice [71]. These genes’ influence on pubertal timing and regulation in the hypothalamus, as well as their impact of sexual differentiation in the rest of the HPG axis could increase our understanding of the mechanisms of development and sex differences.

### Hypothalamic expansion of oligodendrocytes across puberty may be an important precursor to pubertal initiation

Previous studies have reported that oligodendrocyte maturation and myelination can continue into adolescence [124]. However, this expansion has not previously been observed in the hypothalamus specifically. Furthermore, the functional role of oligodendrocytes in regulating hypothalamic hormones involved in pubertal regulation remains an open question [2]. When we integrated our bulk RNA-seq data with publicly available scRNA-seq data [23] in the hypothalamus to estimate cell-type proportions in our samples, we discovered a substantial expansion of OPCs into MOs before and during puberty (Figure 4B, C). Puberty-GWAS genes and oligodendrocyte-mapping age-by-sex-associated genes were overrepresented in oligodendrocyte-pseudotime genes (Figure 5C, D), linking oligodendrocyte development to pubertal regulation in both sexes. Many GWAS hits were enriched in OPCs and are involved in stem-cell differentiation in many cell types [125, 126]. In contrast, age-by-sex and oligodendrocyte-pseudotime associated genes were expressed at all stages of pseudotime expansion and included hormone receptors, core oligodendrocyte markers, and HH disease genes (Figure 5D). For example, *Thra* (thyroid hormone receptor alpha) was primarily expressed in DOs. *Th* also regulated oligodendrocyte expansion in other models [107] and in our bulk RNA-seq, *Th* increases in gene expression before puberty (PD12 < PD22) in female mice. Recently, oligodendrocytes were suggested to be regulated by the HPA axis to facilitate neuroplasticity in major depressive disorder [127], and perhaps a similar phenomena is occurring to facilitate enhanced pubertal neuroplasticity [128]. The identified age-by-sex pattern of *Th* gene expression associated with oligodendrocyte expansion may reflect that oligodendrocytes are highly responsive to sex hormones in a sex-specific manner [129]. For example, progesterone or dihydrotestosterone treated oligodendrocytes increased and decreased proliferation respectively in both male and female mice, however the effect was more extreme in females [129]. Additionally, OPCs show substantial sexual dimorphism in neonatal rats, where female OPCs have stronger transcriptomic responses involved proliferation and migration, while male OPCs have a stronger transcriptomic response involved in differentiation and myelination [130]. Although we identified an enrichment of age-by-sex associated genes mapping to oligodendrocyte development (above and Figures 4, 5), we did not observe sexual dimorphism in oligodendrocyte proportions with RNA-seq deconvolution. Thus, our data lead us to question if proper oligodendrocyte development influences pubertal development by providing the necessary support for the hypothalamic increase in synaptic density and neuronal activation [125, 126] in both sexes.

### Future Directions

Future functional work can be separated into expanding our findings into other tissues, investigating the role of pubertal candidate genes identified in this study, and elucidating whether oligodendrocyte expansion plays an active role in influencing pubertal initiation. Firstly, while puberty is initiated in the hypothalamus, it’s crosstalk between other organs, particularly the pituitary gland and gonads, are key to understanding pubertal regulation overall. We previously investigated sexual dimorphisms across the pubertal transition within the same mice in the pituitary gland and found more evidence of sex-biased gene expression in the pituitary than the hypothalamus, suggesting that the pituitary may have key role in sex differences in puberty [19]. Integrating our current RNA-seq data with that from the pituitary gland and gene expression patterns in the gonads may elucidate regulatory crosstalk between organs [131]. Secondly, investigating differences in transcriptional dynamics, chromatin accessibility, and cell-type composition in mouse models of puberty-relevant candidate genes such as *Hcrt, Oxt*, *Dlk1*, *Cartpt*, or *Gria1* across ages and sexes may elucidate what aspects of pubertal control these candidates may regulate. Lastly, mouse models inhibiting oligodendrocyte expansion could be leveraged to study the pubertal transition further. For example, myelin regulatory factor (*Mrf*) blocks the expansion of OPCs into MOs; however, mice with an *Mrf* knockout die in their third postnatal week, at the same time as we detected OPC expansion in the hypothalamus [132]. Recently, Steadman et al., 2020 developed an inducible *Mrf* knockout that blocks OPC expansion into MOs [133]. Adapting this model to pre-pubertal mice may elucidate if oligodendrocyte expansion influences pubertal initiation. However, this model would not be hypothalamus-specific [133]. To our knowledge, no inducible, hypothalamus specific *Mrf* knockout exists.

### Limitations

One of the limitations of our study is that we investigated the entire hypothalamus rather than sorting the hypothalamus into the paraventricular and arcuate nucleus. Although this is a limitation, the study design allowed us to maximize the number of timepoints and biological replicates while including both sexes. In addition, the scRNA-seq utilized from Kim et al., 2020 contained two timepoints close to puberty, PD14 and PD45 [23]. However, their mice were from two different strains: PD14 mice were CD1 (male and female) and PD45 mice were C57Bl/6J (male) [23]. As such, we integrated these datasets using stringent batch correction [56].

Additionally spatial resolution is important to understand hypothalamic resolution [16, 134], however neither of the bulk RNA-seq or scRNA-seq data in this study is spatially resolved.

## Conclusion

In conclusion, we found that cell type- and sex-aware transcriptomic dynamics in the pubertal hypothalamus are associated with well-established neuropeptide activation and regulation, with the additional highlight of potentially relevant genes including *Hcrt*, *Oxt*, *Dlk1*, *Gria1, and Cartpt*. We discovered that oligodendrocyte expansion occurs in the hypothalamus before and in parallel with early pubertal initiation and that many genes associated with oligodendrocyte expansion relate to pubertal timing and regulation. Our data and interactive Shiny App will allow researchers to visualize the transcriptionally dynamic genes in the hypothalamus and pituitary gland, providing a baseline in postnatal gene expression for the broader scientific community. Together, these data support known mechanisms in pubertal control, provide further insights into sex effects on puberty and suggest novel genes and mechanisms involved in the development of secondary sex characteristics and regulation by the hypothalamus.

## Data availability

RNA-seq data generated for this chapter is available at ArrayExpress (E-MTAB-12340).

## Funding

This work was supported by CIHR grants: 312557 (M.P./M.W./A.G.) and 437197 (Melissa Holmes/M.W./M.P.). M.W. is supported by the Canada Research Chairs Program. R.Q., C.Chan and D.S. were supported in part by NSERC grant RGPIN-2019-07014 to M.W. C.Chan and M.H. were supported by a SickKids RESTRACOMP scholarship. D.S. is supported by NSERC CGS M, PGS D and Ontario Graduate Scholarships. H.H. is supported by the Genome Canada Genomics Technology Platform, The Centre for Applied Genomics. M.M. is supported by NSERC PGS D and the association computing machinery special interest group on high performance computing (ACM/SIGHPC) Intel Computational and Data Science Fellowship. L.U. was supported by the CRS Scholarships for the Next Generation of Scientists.

## Conflict of interest

The authors of this manuscript declare no conflict of interest.

## Notes

https://wilsonlab-sickkids-uoft.shinyapps.io/hypothalamus_gene_shiny/

https://www.ebi.ac.uk/biostudies/arrayexpress/studies/E-MTAB-12340?query=E-MTAB-12340

### Competing Interest Statement

The authors have declared no competing interest.

## Bibliography

1. Day FR, Thompson DJ, Helgason H, Chasman DI, Finucane H, Sulem P, et al. Genomic analyses identify hundreds of variants associated with age at menarche and support a role for puberty timing in cancer risk. Nat Genet. 2017;49:834–841. doi:10.1038/ng.3841.

2. Naulé L, Maione L, Kaiser UB. Puberty, A sensitive window of hypothalamic development and plasticity. Endocrinology. 2021;162. doi:10.1210/endocr/bqaa209.

3. Shin Y-L. An update on the genetic causes of central precocious puberty. Ann Pediatr Endocrinol Metab. 2016;21:66–69. doi:10.6065/apem.2016.21.2.66.

4. Topaloğlu AK. Update on the genetics of idiopathic hypogonadotropic hypogonadism. J Clin Res Pediatr Endocrinol. 2017;9 Suppl 2:113–122. doi:10.4274/jcrpe.2017.S010.

5. Day FR, Bulik-Sullivan B, Hinds DA, Finucane HK, Murabito JM, Tung JY, et al. Shared genetic aetiology of puberty timing between sexes and with health-related outcomes. Nat Commun. 2015;6:8842. doi:10.1038/ncomms9842.

6. Day FR, Elks CE, Murray A, Ong KK, Perry JRB. Puberty timing associated with diabetes, cardiovascular disease and also diverse health outcomes in men and women: the UK Biobank study. Sci Rep. 2015;5:11208. doi:10.1038/srep11208.

7. Sørensen K, Mouritsen A, Aksglaede L, Hagen CP, Mogensen SS, Juul A. Recent secular trends in pubertal timing: implications for evaluation and diagnosis of precocious puberty. Horm Res Paediatr. 2012;77:137–145. doi:10.1159/000336325.

8. Fisher MM, Eugster EA. What is in our environment that effects puberty? Reprod Toxicol. 2014;44:7–14. doi:10.1016/j.reprotox.2013.03.012.

9. Lomniczi A, Wright H, Ojeda SR. Epigenetic regulation of female puberty. Front Neuroendocrinol. 2015;36:90–107. doi:10.1016/j.yfrne.2014.08.003.

10. Hou H, Uusküla-Reimand L, Makarem M, Corre C, Saleh S, Metcalf A, et al. Gene expression profiling of puberty-associated genes reveals abundant tissue and sex-specific changes across postnatal development. Hum Mol Genet. 2017;26:3585–3599. doi:10.1093/hmg/ddx246.

11. Wright H, Aylwin CF, Toro CA, Ojeda SR, Lomniczi A. Polycomb represses a gene network controlling puberty via modulation of histone demethylase Kdm6b expression. Sci Rep. 2021;11:1996. doi:10.1038/s41598-021-81689-4.

12. Lomniczi A, Loche A, Castellano JM, Ronnekleiv OK, Bosch M, Kaidar G, et al. Epigenetic control of female puberty. Nat Neurosci. 2013;16:281–289. doi:10.1038/nn.3319.

13. Abreu AP, Toro CA, Song YB, Navarro VM, Bosch MA, Eren A, et al. MKRN3 inhibits the reproductive axis through actions in kisspeptin-expressing neurons. J Clin Invest. 2020;130:4486–4500. doi:10.1172/JCI136564.

14. Tsutsui K, Son YL, Kiyohara M, Miyata I. Discovery of GnIH and Its Role in Hypothyroidism-Induced Delayed Puberty. Endocrinology. 2018;159:62–68. doi:10.1210/en.2017-00300.

15. Manotas MC, González DM, Céspedes C, Forero C, Rojas Moreno AP. Genetic and epigenetic control of puberty. Sex Dev. 2022;16:1–10. doi:10.1159/000519039.

16. Zhou S, Zang S, Hu Y, Shen Y, Li H, Chen W, et al. Transcriptome-scale spatial gene expression in rat arcuate nucleus during puberty. Cell Biosci. 2022;12:8. doi:10.1186/s13578-022-00745-2.

17. Corre C, Shinoda G, Zhu H, Cousminer DL, Crossman C, Bellissimo C, et al. Sex-specific regulation of weight and puberty by the Lin28/let-7 axis. J Endocrinol. 2016;228:179–191. doi:10.1530/JOE-15-0360.

18. Ong KK, Elks CE, Li S, Zhao JH, Luan J, Andersen LB, et al. Genetic variation in LIN28B is associated with the timing of puberty. Nat Genet. 2009;41:729–733. doi:10.1038/ng.382.

19. Hou H, Chan C, Yuki KE, Sokolowski D, Roy A, Qu R, et al. Postnatal developmental trajectory of sex-biased gene expression in the mouse pituitary gland. Biol Sex Differ. 2022;13:57. doi:10.1186/s13293-022-00467-7.

20. Han X, Burger LL, Garcia-Galiano D, Sim S, Allen SJ, Olson DP, et al. Hypothalamic and Cell-Specific Transcriptomes Unravel a Dynamic Neuropil Remodeling in Leptin-Induced and Typical Pubertal Transition in Female Mice. iScience. 2020;23:101563. doi:10.1016/j.isci.2020.101563.

21. Romanov RA, Zeisel A, Bakker J, Girach F, Hellysaz A, Tomer R, et al. Molecular interrogation of hypothalamic organization reveals distinct dopamine neuronal subtypes. Nat Neurosci. 2017;20:176–188. doi:10.1038/nn.4462.

22. Chen R, Wu X, Jiang L, Zhang Y. Single-Cell RNA-Seq Reveals Hypothalamic Cell Diversity. Cell Rep. 2017;18:3227–3241. doi:10.1016/j.celrep.2017.03.004.

23. Kim DW, Washington PW, Wang ZQ, Lin SH, Sun C, Ismail BT, et al. The cellular and molecular landscape of hypothalamic patterning and differentiation from embryonic to late postnatal development. Nat Commun. 2020;11:4360. doi:10.1038/s41467-020-18231-z.

24. Sokolowski DJ, Faykoo-Martinez M, Erdman L, Hou H, Chan C, Zhu H, et al. Single-cell mapper (scMappR): using scRNA-seq to infer the cell-type specificities of differentially expressed genes. NAR Genom Bioinform. 2021;3:lqab011. doi:10.1093/nargab/lqab011.

25. Chu T, Wang Z, Pe’er D, Danko CG. Cell type and gene expression deconvolution with BayesPrism enables Bayesian integrative analysis across bulk and single-cell RNA sequencing in oncology. Nat Cancer. 2022;3:505–517. doi:10.1038/s43018-022-00356-3.

26. Wang X, Park J, Susztak K, Zhang NR, Li M. Bulk tissue cell type deconvolution with multi-subject single-cell expression reference. Nat Commun. 2019;10:380. doi:10.1038/s41467-018-08023-x.

27. Martin M. Cutadapt removes adapter sequences from high-throughput sequencing reads. EMBnet j. 2011;17:10. doi:10.14806/ej.17.1.200.

28. Dobin A, Davis CA, Schlesinger F, Drenkow J, Zaleski C, Jha S, et al. STAR: ultrafast universal RNA-seq aligner. Bioinformatics. 2013;29:15–21. doi:10.1093/bioinformatics/bts635.

29. Fernandes JD, Zamudio-Hurtado A, Clawson H, Kent WJ, Haussler D, Salama SR, et al. The UCSC repeat browser allows discovery and visualization of evolutionary conflict across repeat families. Mob DNA. 2020;11:13. doi:10.1186/s13100-020-00208-w.

30. Li H, Handsaker B, Wysoker A, Fennell T, Ruan J, Homer N, et al. The Sequence Alignment/Map format and SAMtools. Bioinformatics. 2009;25:2078–2079. doi:10.1093/bioinformatics/btp352.

31. Liao Y, Smyth GK, Shi W. featureCounts: an efficient general purpose program for assigning sequence reads to genomic features. Bioinformatics. 2014;30:923–930. doi:10.1093/bioinformatics/btt656.

32. Harrow J, Frankish A, Gonzalez JM, Tapanari E, Diekhans M, Kokocinski F, et al. GENCODE: the reference human genome annotation for The ENCODE Project. Genome Res. 2012;22:1760–1774. doi:10.1101/gr.135350.111.

33. Frankish A, Diekhans M, Jungreis I, Lagarde J, Loveland JE, Mudge JM, et al. GENCODE 2021. Nucleic Acids Res. 2021;49:D916–D923. doi:10.1093/nar/gkaa1087.

34. Risso D, Ngai J, Speed TP, Dudoit S. Normalization of RNA-seq data using factor analysis of control genes or samples. Nat Biotechnol. 2014;32:896–902. doi:10.1038/nbt.2931.

35. Gu Z, Eils R, Schlesner M. Complex heatmaps reveal patterns and correlations in multidimensional genomic data. Bioinformatics. 2016;32:2847–2849. doi:10.1093/bioinformatics/btw313.

36. Wold S, Esbensen K, Geladi P. Principal component analysis. Chemometrics and Intelligent Laboratory Systems. 1987;2:37–52. doi:10.1016/0169-7439(87)80084-9.

37. Wickham H. ggplot2: Elegant graphics for data analysis. New York: Springer; 2009.

38. Love MI, Huber W, Anders S. Moderated estimation of fold change and dispersion for RNA-seq data with DESeq2. Genome Biol. 2014;15:550. doi:10.1186/s13059-014-0550-8.

39. Kaiser HF. The varimax criterion for analytic rotation in factor analysis. Psychometrika. 1958;23:187–200. doi:10.1007/BF02289233.

40. Cureton EE, Mulaik SA. The weighted varimax rotation and the promax rotation. Psychometrika. 1975;40:183–195. doi:10.1007/BF02291565.

41. Borchers HW. Practical Numerical Math Functions [R package pracma version 2.3.3]. 2021.

42. Korthauer K, Kimes PK, Duvallet C, Reyes A, Subramanian A, Teng M, et al. A practical guide to methods controlling false discoveries in computational biology. Genome Biol. 2019;20:118. doi:10.1186/s13059-019-1716-1.

43. Paczkowska M, Barenboim J, Sintupisut N, Fox NS, Zhu H, Abd-Rabbo D, et al. Integrative pathway enrichment analysis of multivariate omics data. Nat Commun. 2020;11:735. doi:10.1038/s41467-019-13983-9.

44. Reimand J, Isserlin R, Voisin V, Kucera M, Tannus-Lopes C, Rostamianfar A, et al. Pathway enrichment analysis and visualization of omics data using g:Profiler, GSEA, Cytoscape and EnrichmentMap. Nat Protoc. 2019;14:482–517. doi:10.1038/s41596-018-0103-9.

45 . Reimand J, Arak T, Adler P, Kolberg L, Reisberg S, Peterson H, et al. g:Profiler-a web server for functional interpretation of gene lists (2016 update). Nucleic Acids Res. 2016;44:W83–9. doi:10.1093/nar/gkw199.

46. Sokolowski DJ, Ahn J, Erdman L, Hou H, Ellis K, Wang L, et al. Differential Expression Enrichment Tool (DEET): an interactive atlas of human differential gene expression. NAR Genom Bioinform. 2023;5:lqad003. doi:10.1093/nargab/lqad003.

47. Tomczak K, Czerwińska P. The Cancer Genome Atlas (TCGA): an immeasurable source of knowledge. Contemp Oncol (Pozn). 2015;19:A68–77. doi:10.5114/wo.2014.47136.

48. GTEx Consortium. Human genomics. The Genotype-Tissue Expression (GTEx) pilot analysis: multitissue gene regulation in humans. Science. 2015;348:648–660. doi:10.1126/science.1262110.

49. Kodama Y, Shumway M, Leinonen R, International Nucleotide Sequence Database Collaboration. The Sequence Read Archive: explosive growth of sequencing data. Nucleic Acids Res. 2012;40 Database issue:D54–6. doi:10.1093/nar/gkr854.

50. Collado-Torres L, Nellore A, Kammers K, Ellis SE, Taub MA, Hansen KD, et al. Reproducible RNA-seq analysis using recount2. Nat Biotechnol. 2017;35:319–321. doi:10.1038/nbt.3838.

51. Allison MB, Patterson CM, Krashes MJ, Lowell BB, Myers MG, Olson DP. TRAP-seq defines markers for novel populations of hypothalamic and brainstem LepRb neurons. Mol Metab. 2015;4:299–309. doi:10.1016/j.molmet.2015.01.012.

52. Hafemeister C, Satija R. Normalization and variance stabilization of single-cell RNA-seq data using regularized negative binomial regression. Genome Biol. 2019;20:296. doi:10.1186/s13059-019-1874-1.

53. Hao Y, Hao S, Andersen-Nissen E, Mauck WM, Zheng S, Butler A, et al. Integrated analysis of multimodal single-cell data. Cell. 2021;184:3573–3587. doi:10.1016/j.cell.2021.04.048.

54. Zhang X, Lan Y, Xu J, Quan F, Zhao E, Deng C, et al. CellMarker: a manually curated resource of cell markers in human and mouse. Nucleic Acids Res. 2019;47:D721–D728. doi:10.1093/nar/gky900.

55. Franzén O, Gan L-M, Björkegren JLM. PanglaoDB: a web server for exploration of mouse and human single-cell RNA sequencing data. Database (Oxford). 2019;2019. doi:10.1093/database/baz046.

56. Stuart T, Butler A, Hoffman P, Hafemeister C, Papalexi E, Mauck WM, et al. Comprehensive Integration of Single-Cell Data. Cell. 2019;177:1888–1902.e21. doi:10.1016/j.cell.2019.05.031.

57. Luecken MD, Büttner M, Chaichoompu K, Danese A, Interlandi M, Mueller MF, et al. Benchmarking atlas-level data integration in single-cell genomics. Nat Methods. 2022;19:41–50. doi:10.1038/s41592-021-01336-8.

58. Upton GJG. Fisher’s Exact Test. Journal of the Royal Statistical Society Series A (Statistics in Society). 1992;155:395. doi:10.2307/2982890.

59. Finak G, McDavid A, Yajima M, Deng J, Gersuk V, Shalek AK, et al. MAST: a flexible statistical framework for assessing transcriptional changes and characterizing heterogeneity in single-cell RNA sequencing data. Genome Biol. 2015;16:278. doi:10.1186/s13059-015-0844-5.

60. Gong T, Szustakowski JD. DeconRNASeq: a statistical framework for deconvolution of heterogeneous tissue samples based on mRNA-Seq data. Bioinformatics. 2013;29:1083–1085. doi:10.1093/bioinformatics/btt090.

61. Altboum Z, Steuerman Y, David E, Barnett-Itzhaki Z, Valadarsky L, Keren-Shaul H, et al. Digital cell quantification identifies global immune cell dynamics during influenza infection. Mol Syst Biol. 2014;10:720. doi:10.1002/msb.134947.

62. Langfelder P, Horvath S. WGCNA: an R package for weighted correlation network analysis. BMC Bioinformatics. 2008;9:559. doi:10.1186/1471-2105-9-559.

63. Chen B, Khodadoust MS, Liu CL, Newman AM, Alizadeh AA. Profiling Tumor Infiltrating Immune Cells with CIBERSORT. Methods Mol Biol. 2018;1711:243–259. doi:10.1007/978-1-4939-7493-1_12.

64. Newman AM, Steen CB, Liu CL, Gentles AJ, Chaudhuri AA, Scherer F, et al. Determining cell type abundance and expression from bulk tissues with digital cytometry. Nat Biotechnol. 2019;37:773–782. doi:10.1038/s41587-019-0114-2.

65. Frishberg A, Peshes-Yaloz N, Cohn O, Rosentul D, Steuerman Y, Valadarsky L, et al. Cell composition analysis of bulk genomics using single-cell data. Nat Methods. 2019;16:327–332. doi:10.1038/s41592-019-0355-5.

66. Danziger SA, Gibbs DL, Shmulevich I, McConnell M, Trotter MWB, Schmitz F, et al. ADAPTS: Automated deconvolution augmentation of profiles for tissue specific cells. PLoS One. 2019;14:e0224693. doi:10.1371/journal.pone.0224693.

67. Street K, Risso D, Fletcher RB, Das D, Ngai J, Yosef N, et al. Slingshot: cell lineage and pseudotime inference for single-cell transcriptomics. BMC Genomics. 2018;19:477. doi:10.1186/s12864-018-4772-0.

68. Van den Berge K, Roux de Bézieux H, Street K, Saelens W, Cannoodt R, Saeys Y, et al. Trajectory-based differential expression analysis for single-cell sequencing data. Nat Commun. 2020;11:1201. doi:10.1038/s41467-020-14766-3.

69. ENCODE Project Consortium. The ENCODE (encyclopedia of DNA elements) project. Science. 2004;306:636–640. doi:10.1126/science.1105136.

70. Valadares LP, Meireles CG, De Toledo IP, Santarem de Oliveira R, Gonçalves de Castro LC, Abreu AP, et al. MKRN3 Mutations in Central Precocious Puberty: A Systematic Review and Meta-Analysis. J Endocr Soc. 2019;3:979–995. doi:10.1210/js.2019-00041.

71. Venancio JC, Margatho LO, Rorato R, Rosales RRC, Debarba LK, Coletti R, et al. Short-Term High-Fat Diet Increases Leptin Activation of CART Neurons and Advances Puberty in Female Mice. Endocrinology. 2017;158:3929–3942. doi:10.1210/en.2017-00452.

72. Macedo DB, Abreu AP, Tellez SL, Naule L, Kim HK, Capo-Battaglia A, et al. SUN-100 Mice Lacking Paternally Expressed DLK1 Reach Puberty at a Lower Body Weight Than Littermate Controls. Journal of the Endocrine Society. 2020;4. doi:10.1210/jendso/bvaa046.1567.

73. Wiemann JN, Clifton DK, Steiner RA. Pubertal changes in gonadotropin-releasing hormone and proopiomelanocortin gene expression in the brain of the male rat. Endocrinology. 1989;124:1760–1767. doi:10.1210/endo-124-4-1760.

74. Marques S, Zeisel A, Codeluppi S, van Bruggen D, Mendanha Falcão A, Xiao L, et al. Oligodendrocyte heterogeneity in the mouse juvenile and adult central nervous system. Science. 2016;352:1326–1329. doi:10.1126/science.aaf6463.

75. Chuang N, Mori S, Yamamoto A, Jiang H, Ye X, Xu X, et al. An MRI-based atlas and database of the developing mouse brain. Neuroimage. 2011;54:80–89. doi:10.1016/j.neuroimage.2010.07.043.

76. León S, Fergani C, Talbi R, Simavli S, Maguire CA, Gerutshang A, et al. Characterization of the role of NKA in the control of puberty onset and gonadotropin release in the female mouse. Endocrinology. 2019;160:2453–2463. doi:10.1210/en.2019-00195.

77. Nurhidayat, Tsukamoto Y, Sigit K, Sasaki F. Sex differentiation of growth hormone-releasing hormone and somatostatin neurons in the mouse hypothalamus: an immunohistochemical and morphological study. Brain Res. 1999;821:309–321. doi:10.1016/s0006-8993(99)01081-1.

78. Fonseca DJ, Ojeda D, Lakhal B, Braham R, Eggers S, Turbitt E, et al. CITED2 mutations potentially cause idiopathic premature ovarian failure. Transl Res. 2012;160:384–388. doi:10.1016/j.trsl.2012.05.006.

79. Tsai P-S, Brooks LR, Rochester JR, Kavanaugh SI, Chung WCJ. Fibroblast growth factor signaling in the developing neuroendocrine hypothalamus. Front Neuroendocrinol. 2011;32:95–107. doi:10.1016/j.yfrne.2010.11.002.

80. Palijan A, Fernandes I, Verway M, Kourelis M, Bastien Y, Tavera-Mendoza LE, et al. Ligand-dependent corepressor LCoR is an attenuator of progesterone-regulated gene expression. J Biol Chem. 2009;284:30275–30287. doi:10.1074/jbc.M109.051201.

81. Curtin D, Jenkins S, Farmer N, Anderson AC, Haisenleder DJ, Rissman E, et al. Androgen suppression of GnRH-stimulated rat LHbeta gene transcription occurs through Sp1 sites in the distal GnRH-responsive promoter region. Mol Endocrinol. 2001;15:1906–1917. doi:10.1210/mend.15.11.0723.

82. Egan OK, Inglis MA, Anderson GM. Leptin signaling in agrp neurons modulates puberty onset and adult fertility in mice. J Neurosci. 2017;37:3875–3886. doi:10.1523/JNEUROSCI.3138-16.2017.

83. Giacobini P, Wray S. Cholecystokinin directly inhibits neuronal activity of primary gonadotropin-releasing hormone cells through cholecystokinin-1 receptor. Endocrinology. 2007;148:63–71. doi:10.1210/en.2006-0758.

84. Mul JD, Yi C-X, van den Berg SAA, Ruiter M, Toonen PW, van der Elst MCJ, et al. Pmch expression during early development is critical for normal energy homeostasis. Am J Physiol Endocrinol Metab. 2010;298:E477–88. doi:10.1152/ajpendo.00154.2009.

85. Tao Y-H, Sharif N, Zeng B-H, Cai Y-Y, Guo Y-X. Lateral ventricle injection of orexin-A ameliorates central precocious puberty in rat via inhibiting the expression of MEG3. Int J Clin Exp Pathol. 2015;8:12564–12570.

86. Poli F, Pizza F, Mignot E, Ferri R, Pagotto U, Taheri S, et al. High prevalence of precocious puberty and obesity in childhood narcolepsy with cataplexy. Sleep. 2013;36:175–181. doi:10.5665/sleep.2366.

87. Parent A-S, Rasier G, Matagne V, Lomniczi A, Lebrethon M-C, Gérard A, et al. Oxytocin facilitates female sexual maturation through a glia-to-neuron signaling pathway. Endocrinology. 2008;149:1358–1365. doi:10.1210/en.2007-1054.

88. Salian-Mehta S, Xu M, Knox AJ, Plummer L, Slavov D, Taylor M, et al. Functional consequences of AXL sequence variants in hypogonadotropic hypogonadism. J Clin Endocrinol Metab. 2014;99:1452–1460. doi:10.1210/jc.2013-3426.

89. Lappalainen S, Voutilainen R, Utriainen P, Laakso M, Jääskeläinen J. Genetic variation of FTO and TCF7L2 in premature adrenarche. Metab Clin Exp. 2009;58:1263–1269. doi:10.1016/j.metabol.2009.03.025.

90. Leinonen R, Sugawara H, Shumway M, International Nucleotide Sequence Database Collaboration. The sequence read archive. Nucleic Acids Res. 2011;39 Database issue:D19–21. doi:10.1093/nar/gkq1019.

91. Bao Z-S, Chen H-M, Yang M-Y, Zhang C-B, Yu K, Ye W-L, et al. RNA-seq of 272 gliomas revealed a novel, recurrent PTPRZ1-MET fusion transcript in secondary glioblastomas. Genome Res. 2014;24:1765–1773. doi:10.1101/gr.165126.113.

92. He Z, Bammann H, Han D, Xie G, Khaitovich P. Conserved expression of lincRNA during human and macaque prefrontal cortex development and maturation. RNA. 2014;20:1103–1111. doi:10.1261/rna.043075.113.

93. Ladouceur CD, Peper JS, Crone EA, Dahl RE. White matter development in adolescence: the influence of puberty and implications for affective disorders. Dev Cogn Neurosci. 2012;2:36–54. doi:10.1016/j.dcn.2011.06.002.

94. Su Y, Huang J, Zhao X, Lu H, Wang W, Yang XO, et al. Interleukin-17 receptor D constitutes an alternative receptor for interleukin-17A important in psoriasis-like skin inflammation. Sci Immunol. 2019;4. doi:10.1126/sciimmunol.aau9657.

95. Koncina E, Roth L, Gonthier B, Bagnard D. Role of semaphorins during axon growth and guidance. Adv Exp Med Biol. 2007;621:50–64. doi:10.1007/978-0-387-76715-4_4.

96. Oishi H, Sasaki T, Nagano F, Ikeda W, Ohya T, Wada M, et al. Localization of the Rab3 small G protein regulators in nerve terminals and their involvement in Ca2+-dependent exocytosis. J Biol Chem. 1998;273:34580–34585. doi:10.1074/jbc.273.51.34580.

97. Mooney SJ, Douglas NR, Holmes MM. Peripheral administration of oxytocin increases social affiliation in the naked mole-rat (Heterocephalus glaber). Horm Behav. 2014;65:380–385. doi:10.1016/j.yhbeh.2014.02.003.

98. Anttila R, Siimes MA. Serum transferrin and ferritin in pubertal boys: relations to body growth, pubertal stage, erythropoiesis, and iron deficiency. Am J Clin Nutr. 1996;63:179–183. doi:10.1093/ajcn/63.2.179.

99. Sutton GJ, Poppe D, Simmons RK, Walsh K, Nawaz U, Lister R, et al. Comprehensive evaluation of deconvolution methods for human brain gene expression. Nat Commun. 2022;13:1358. doi:10.1038/s41467-022-28655-4.

100. Connor JR, Menzies SL. Relationship of iron to oligodendrocytes and myelination. Glia. 1996;17:83–93. doi:10.1002/(SICI)1098-136(199606)17:2<83::AID-GLIA1>3.0.CO;2-7.

101. Macedo DB, Kaiser UB. DLK1, notch signaling and the timing of puberty. Semin Reprod Med. 2019;37:174–181. doi:10.1055/s-0039-3400963.

102. Sugimoto M, Sasaki S, Watanabe T, Nishimura S, Ideta A, Yamazaki M, et al. Ionotropic glutamate receptor AMPA 1 is associated with ovulation rate. PLoS One. 2010;5:e13817. doi:10.1371/journal.pone.0013817.

103. Sanchez-Garrido MA, Tena-Sempere M. Metabolic control of puberty: roles of leptin and kisspeptins. Horm Behav. 2013;64:187–194. doi:10.1016/j.yhbeh.2013.01.014.

104. Donato J, Cravo RM, Frazão R, Gautron L, Scott MM, Lachey J, et al. Leptin’s effect on puberty in mice is relayed by the ventral premammillary nucleus and does not require signaling in Kiss1 neurons. J Clin Invest. 2011;121:355–368. doi:10.1172/JCI45106.

105. Dauber A, Cunha-Silva M, Macedo DB, Brito VN, Abreu AP, Roberts SA, et al. Paternally inherited DLK1 deletion associated with familial central precocious puberty. J Clin Endocrinol Metab. 2017;102:1557–1567. doi:10.1210/jc.2016-3677.

106. Noda Y, Ota K, Shirasawa T, Shimizu T. Copper/zinc superoxide dismutase insufficiency impairs progesterone secretion and fertility in female mice. Biol Reprod. 2012;86:1–8. doi:10.1095/biolreprod.111.092999.

107. Rodríguez-Peña A. Oligodendrocyte development and thyroid hormone. J Neurobiol. 1999;40:497–512. doi:10.1002/(sici)1097-4695(19990915)40:4<497::aid-neu7>3.0.co;2-#.

108. Schaefer T, Lengerke C. SOX2 protein biochemistry in stemness, reprogramming, and cancer: the PI3K/AKT/SOX2 axis and beyond. Oncogene. 2020;39:278–292. doi:10.1038/s41388-019-0997-x.

109. Zhang S, Zhu X, Gui X, Croteau C, Song L, Xu J, et al. Sox2 Is Essential for Oligodendroglial Proliferation and Differentiation during Postnatal Brain Myelination and CNS Remyelination. J Neurosci. 2018;38:1802–1820. doi:10.1523/JNEUROSCI.1291-17.2018.

110. Kelberman D, Rizzoti K, Avilion A, Bitner-Glindzicz M, Cianfarani S, Collins J, et al. Mutations within Sox2/SOX2 are associated with abnormalities in the hypothalamo-pituitary-gonadal axis in mice and humans. J Clin Invest. 2006;116:2442–2455. doi:10.1172/JCI28658.

111. White MD, Angiolini JF, Alvarez YD, Kaur G, Zhao ZW, Mocskos E, et al. Long-Lived Binding of Sox2 to DNA Predicts Cell Fate in the Four-Cell Mouse Embryo. Cell. 2016;165:75–87. doi:10.1016/j.cell.2016.02.032.

112. Kelberman D, de Castro SCP, Huang S, Crolla JA, Palmer R, Gregory JW, et al. SOX2 plays a critical role in the pituitary, forebrain, and eye during human embryonic development. J Clin Endocrinol Metab. 2008;93:1865–1873. doi:10.1210/jc.2007-2337.

113. Goldman SA, Kuypers NJ. How to make an oligodendrocyte. Development. 2015;142:3983–3995. doi:10.1242/dev.126409.

114. He D, Marie C, Zhao C, Kim B, Wang J, Deng Y, et al. Chd7 cooperates with Sox10 and regulates the onset of CNS myelination and remyelination. Nat Neurosci. 2016;19:678–689. doi:10.1038/nn.4258.

115. Muthusamy K, Sudhakar SV, Yoganathan S, Thomas MM, Alexander M. Hypomyelination, hypodontia, hypogonadotropic hypogonadism (4H) syndrome with vertebral anomalies: A novel association. J Child Neurol. 2015;30:937–941. doi:10.1177/0883073814541470.

116. Mahler B, Kamperis K, Ankarberg-Lindgren C, Frøkiær J, Djurhuus JC, Rittig S. Puberty alters renal water handling. Am J Physiol Renal Physiol. 2013;305:F1728–35. doi:10.1152/ajprenal.00283.2013.

117. Melzi S, Prevot V, Peyron C. Precocious puberty in narcolepsy type 1: Orexin loss and/or neuroinflammation, which is to blame? Sleep Med Rev. 2022;65:101683. doi:10.1016/j.smrv.2022.101683.

118. Gaskins GT, Moenter SM. Orexin a suppresses gonadotropin-releasing hormone (GnRH) neuron activity in the mouse. Endocrinology. 2012;153:3850–3860. doi:10.1210/en.2012-1300.

119. Hosseini A, Khazali H. Central orexin A affects reproductive axis by modulation of hypothalamic kisspeptin/neurokinin b/dynorphin secreting neurons in the male wistar rats. Neuromolecul Med. 2018;20:525–536. doi:10.1007/s12017-018-8506-x.

120. Lucien JN, Ortega MT, Shaw ND. Sleep and Puberty. Current Opinion in Endocrine and Metabolic Research. 2021;17:1–7. doi:10.1016/j.coemr.2020.09.009.

121. Miller TV, Caldwell HK. Oxytocin during Development: Possible Organizational Effects on Behavior. Front Endocrinol (Lausanne). 2015;6:76. doi:10.3389/fendo.2015.00076.

122. Boyle EA, Li YI, Pritchard JK. An expanded view of complex traits: from polygenic to omnigenic. Cell. 2017;169:1177–1186. doi:10.1016/j.cell.2017.05.038.

123. Montenegro L, Labarta JI, Piovesan M, Canton APM, Corripio R, Soriano-Guillén L, et al. Novel Genetic and Biochemical Findings of DLK1 in Children with Central Precocious Puberty: A Brazilian-Spanish Study. J Clin Endocrinol Metab. 2020;105. doi:10.1210/clinem/dgaa461.

124. Williamson JM, Lyons DA. Myelin Dynamics Throughout Life: An Ever-Changing Landscape? Front Cell Neurosci. 2018;12:424. doi:10.3389/fncel.2018.00424.

125. Ojeda SR, Lomniczi A, Sandau U. Contribution of glial-neuronal interactions to the neuroendocrine control of female puberty. Eur J Neurosci. 2010;32:2003–2010. doi:10.1111/j.1460-9568.2010.07515.x.

126. Clasadonte J, Prevot V. The special relationship: glia-neuron interactions in the neuroendocrine hypothalamus. Nat Rev Endocrinol. 2018;14:25–44. doi:10.1038/nrendo.2017.124.

127. Zhou B, Zhu Z, Ransom BR, Tong X. Oligodendrocyte lineage cells and depression. Mol Psychiatry. 2021;26:103–117. doi:10.1038/s41380-020-00930-0.

128. Laube C, van den Bos W, Fandakova Y. The relationship between pubertal hormones and brain plasticity: Implications for cognitive training in adolescence. Dev Cogn Neurosci. 2020;42:100753. doi:10.1016/j.dcn.2020.100753.

129. Swamydas M, Bessert D, Skoff R. Sexual dimorphism of oligodendrocytes is mediated by differential regulation of signaling pathways. J Neurosci Res. 2009;87:3306–3319. doi:10.1002/jnr.21943.

130. Yasuda K, Maki T, Kinoshita H, Kaji S, Toyokawa M, Nishigori R, et al. Sex-specific differences in transcriptomic profiles and cellular characteristics of oligodendrocyte precursor cells. Stem Cell Res. 2020;46:101866. doi:10.1016/j.scr.2020.101866.

131. Mack KL, Phifer-Rixey M, Harr B, Nachman MW. Gene Expression Networks Across Multiple Tissues Are Associated with Rates of Molecular Evolution in Wild House Mice. Genes (Basel). 2019;10. doi:10.3390/genes10030225.

132. Emery B, Agalliu D, Cahoy JD, Watkins TA, Dugas JC, Mulinyawe SB, et al. Myelin gene regulatory factor is a critical transcriptional regulator required for CNS myelination. Cell. 2009;138:172–185. doi:10.1016/j.cell.2009.04.031.

133. Steadman PE, Xia F, Ahmed M, Mocle AJ, Penning ARA, Geraghty AC, et al. Disruption of oligodendrogenesis impairs memory consolidation in adult mice. Neuron. 2020;105:150–164.e6. doi:10.1016/j.neuron.2019.10.013.

134. Moffitt JR, Bambah-Mukku D, Eichhorn SW, Vaughn E, Shekhar K, Perez JD, et al. Molecular, spatial, and functional single-cell profiling of the hypothalamic preoptic region. Science. 2018;362. doi:10.1126/science.aau5324.

