## Supplementary Figures and Tables for "Age, sex, and cell type-resolved hypothalamic gene expression across the pubertal transition in mice"

(7) CIFAR, Toronto, ON, Canada

(8) Division of Endocrinology, The Hospital for Sick Children, Toronto, ON, Canada

(9) Institute of Medical Science, University of Toronto, Toronto, ON, Canada

(10) Departments of Pediatrics and Physiology, University of Toronto, Toronto, ON, Canada

- CIFAR and MARS for Anna Goldenberg

*corresponding author

### Table of Contents

**Supplementary Figure S1**. Quality control metrics of our high-throughput mRNA-seq data. (Page 4).

**Supplementary Figure S2**. Summary of differentially expressed genes (DEGs) between PD32 vs. PD37 in female mice. (Page 5).

**Supplementary Figure S3**. Summary of transcriptional co-repressors that decrease in expression during puberty and pubertal-relevant neuropeptides that increase in expression during puberty. (Page 7).

**Supplementary Figure S4**. Overview of sex differences across each pubertal timepoint. A) Volcano plots of sex differences at each time point. (Page 9).

**Supplementary Figure S5**. Differentially expressed genes (DEGs) between PD12 vs. PD22 male and female mice enriched against 3162 human differential gene expression comparisons using the Differential Expression Enrichment Tool (DEET) and ActivePathways. (Page 11).

**Supplementary Figure S6**. Correlation plots between PD12 vs. PD22 males and the human comparisons within the DEET database that contain the top 3 most correlated DEGs. (Page 14).

**Supplementary Figure S7**. Summary of differentially expressed hypogonadotropic hypogonadism genes and puberty genome-wide association study (GWAS) genes. (Page 16).

**Supplementary Figure S8**. Enrichment of age-by-sex associated genes against 3162 human RNA-seq comparisons stored in the DEET dataset. (Page 18).

**Supplementary Figure S9**. Distribution of cell-type proportions measured from MuSiC-NNLS in cell-types that were predicted to have >3% of the total sample. (Page 20).

**Supplementary Figure S10**. Heatmap of gene-normalized cell-weighted fold-changes (cwFold-changes) of the 129 age-by-sex associated genes and are DE in the complementary direction in the scRNA-seq data. (Page 22).

**Supplementary Figure S11**. GeneMANIA plot of the 21 neuron- neuroendocrine-mapping age-by-sex associated genes that are detected as translated in hypothalamic LepRb+ neurons in Trap-seq from Allison et al., 2015. (Page 24).

**Supplementary Figure S12**. Pseudotime of hypothalamic oligodendrocyte development. Heatmap of RNA polymerase subunit genes associated with pseudotime. (Page 24).

**Supplementary Table S1**. Summary of RNA-seq sample quality and read mapping using Qualimap (Page 26).

**Supplementary Table S2**. Correlation and number of cell-types detected (>1% of the total population) between cell-type proportions from RNA-seq deconvolution and cell-type proportions from scRNA-seq data in the mouse hypothalamus (Page 27).

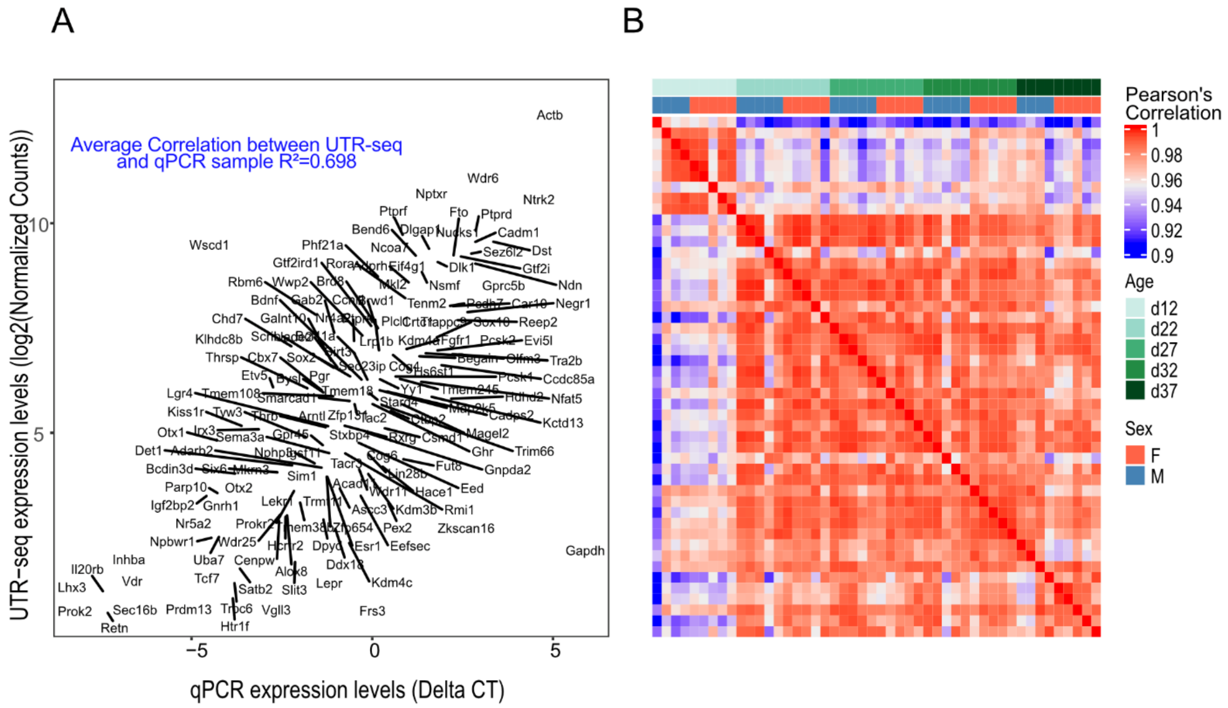


**Supplementary Figure S1. Quality control metrics of our high-throughput mRNA-seq data.** A) Correlation between UTR-seq and qPCR of 182 genes deriving from the same RNA. The X-axis represents each gene's average qPCR expression level across all 48 samples. The Y-axis is the average log2(RUV-seq)-normalized gene expression profiles across all 48 samples. B) Correlation heatmap of all UTR-seq samples, where all genes were included. Rows and columns are samples, and the heatmap is populated by Pearson’s correlation between them. Columns are annotated by age and sex and labeled by replicate number.

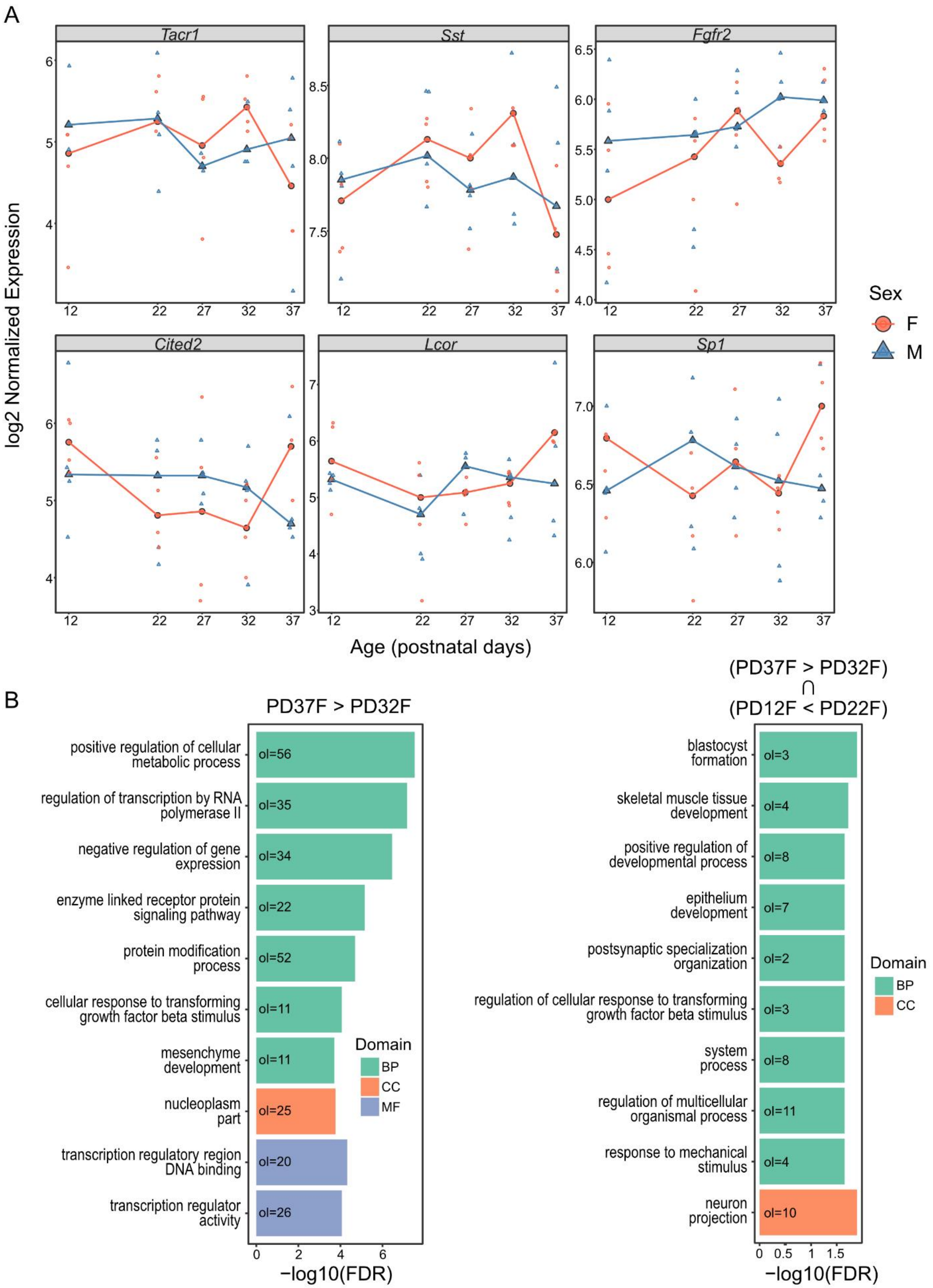
A
Tacr1
Sst
Fgfr2
Cited2
Lcor
Sp1
log2 Normalized Expression
Age (postnatal days)
Sex
F
M
B
PD37F > PD32F
positive regulation of cellular metabolic process
regulation of transcription by RNA polymerase II
negative regulation of gene expression
enzyme linked receptor protein signaling pathway
protein modification process
cellular response to transforming growth factor beta stimulus
mesenchyme development
nucleoplasm part
transcription regulatory region DNA binding
transcription regulator activity
ol=56
ol=35
ol=34
ol=22
ol=52
ol=11
ol=11
ol=25
ol=20
ol=26
Domain
BP
CC
MF
−log10(FDR)
(PD37F > PD32F) ∩ (PD12F < PD22F)
blastocyst formation
skeletal muscle tissue development
positive regulation of developmental process
epithelium development
postsynaptic specialization organization
regulation of cellular response to transforming growth factor beta stimulus
system process
regulation of multicellular organismal process
response to mechanical stimulus
neuron projection
ol=3
ol=4
ol=8
ol=7
ol=2
ol=3
ol=8
ol=11
ol=4
ol=10
Domain
BP
CC
−log10(FDR)

**Supplementary Figure S2. Summary of differentially expressed genes (DEGs) between PD32 vs. PD37 in female mice.** A) Distribution of normalized counts from hormone regulators/producers (*Tacr1*, *Sst*) and transcriptional regulators (*Fgfr2*, *Cited2*, *Lcor*, *Sp1*). The X-axis is age, and the Y-axis is log2-transformed RUVseq and ERCC-spike in normalized counts. Red lines and circles represent female samples, while blue lines and triangles represent male samples. B) Barplot of pathway enrichment of upregulated DEGs between PD37 vs. PD32 in female mice. C) Barplot of pathway enrichment of the overlap of upregulated DEGs between PD37 vs. PD32 in female mice and of downregulated DEGs between PD12 vs. PD22 in female mice.

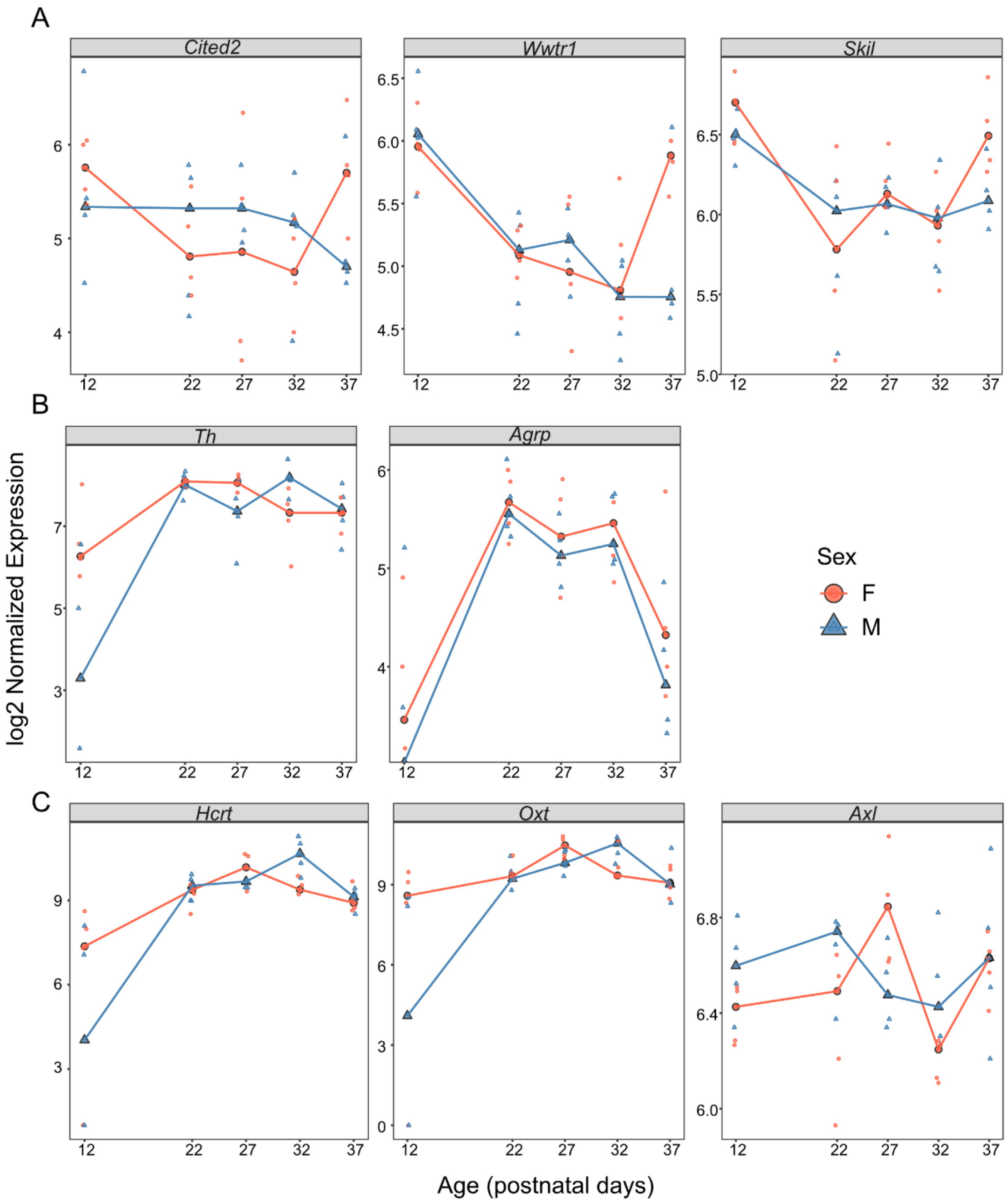


**Supplementary Figure S3. Summary of transcriptional co-repressors that decrease in expression during puberty and pubertal-relevant neuropeptides that increase in expression during puberty.** The X-axis is age, and the Y-axis is log2-transformed RUVseq and ERCC-spike in normalized counts. Red lines and circles represent female samples, while blue lines and triangles represent male samples. A) Transcriptional co-repressors that increase in expression between PD12 vs. PD22 in females and increase in expression between PD32 vs. PD37 in females. B) Direct pubertal regulators that increase in gene expression between PD12 vs. PD22 in females. C) Hormonal neuropeptide genes which peak in expression at PD27 in females, when half of the female mice have undergone vaginal opening.

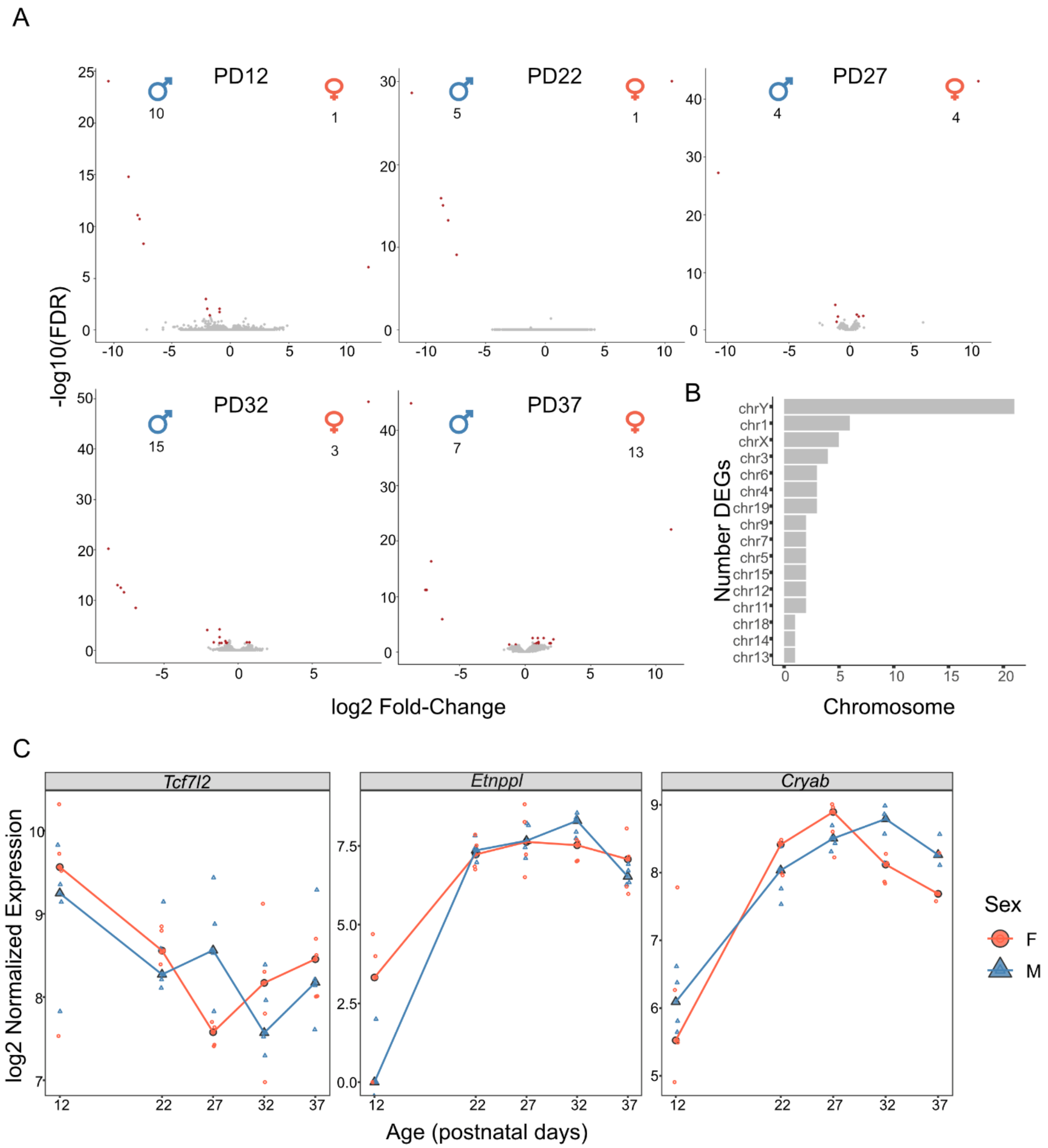


**Supplementary Figure S4. Overview of sex differences across each pubertal timepoint. A) Volcano plots of sex differences at each time point.** The X-axis is the fold-change of each gene, and the Y-axis is the -log10(FDR-adjusted p-value). Genes with positive fold-changes are female-biased, and genes with negative fold-changes are male-biased. B) Barplot of the chromosomal distribution of sex differences at any timepoint. C) Gene expression distribution of three sex-biased genes that were previously associated with puberty. The X-axis is age, and the Y-axis is log2-transformed RUVseq and ERCC-spike in normalized counts. Red lines and circles represent female samples, while blue lines and triangles represent male samples.

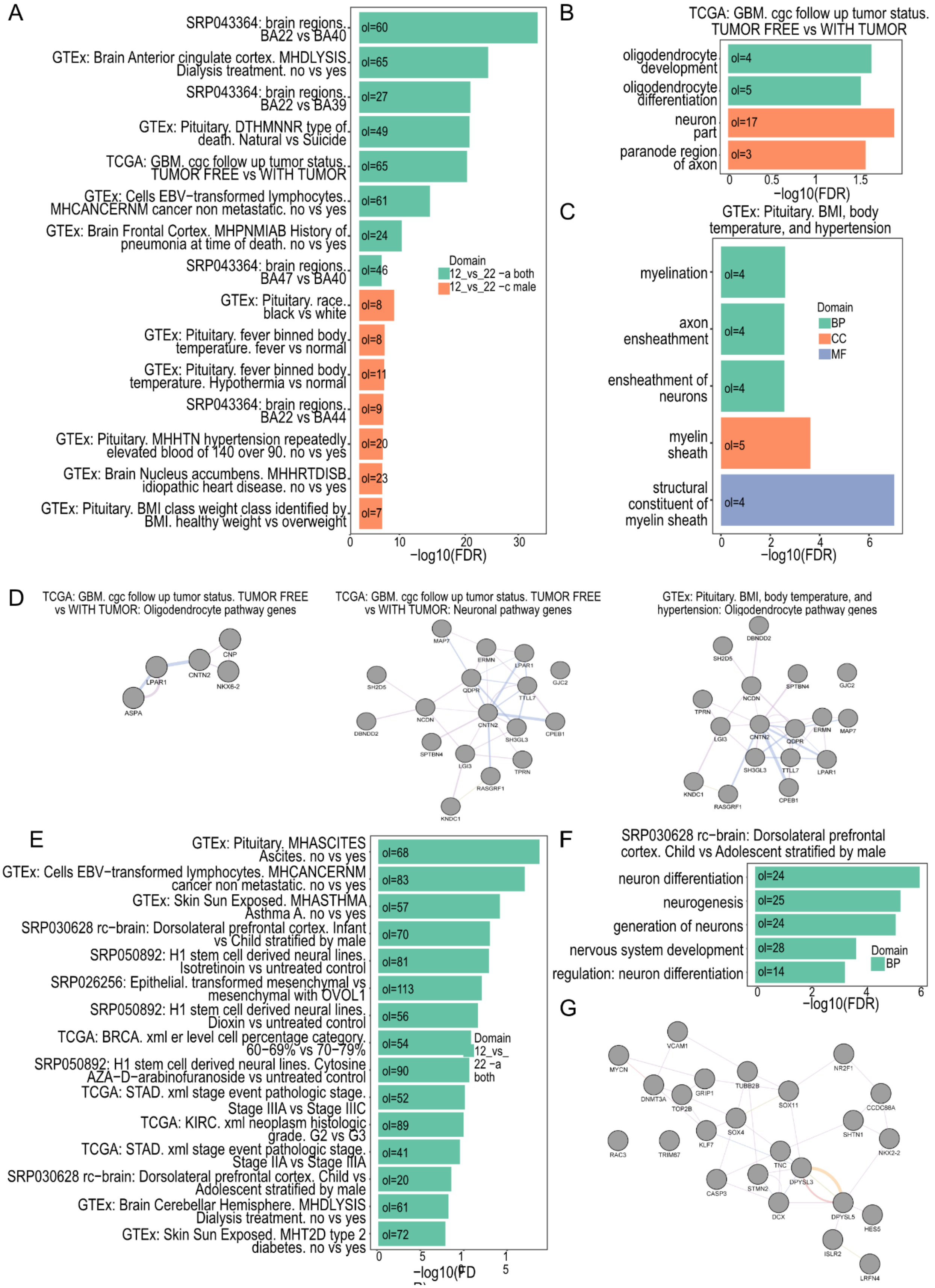

A
SRP043364: brain regions. BA22 vs BA40
GTEx: Brain Anterior cingulate cortex. MHDLYSIS Dialysis treatment. no vs yes
SRP043364: brain regions. BA22 vs BA39
GTEx: Pituitary. DTHMNNR type of death. Natural vs Suicide
TCGA: GBM. cgc follow up tumor status. TUMOR FREE vs WITH TUMOR
GTEx: Cells EBV−transformed lymphocytes. MHCANCERNM cancer non metastatic. no vs yes
GTEx: Brain Frontal Cortex. MHPNMIAB History of pneumonia at time of death. no vs yes
SRP043364: brain regions. BA47 vs BA40
GTEx: Pituitary. race. black vs white
GTEx: Pituitary. fever binned body temperature. fever vs normal
GTEx: Pituitary. fever binned body temperature. Hypothermia vs normal
SRP043364: brain regions. BA22 vs BA44
GTEx: Pituitary. MHHTN hypertension repeatedly elevated blood of 140 over 90. no vs yes
GTEx: Brain Nucleus accumbens. MHHRTDISB idiopathic heart disease. no vs yes
GTEx: Pituitary. BMI class weight class identified by BMI. healthy weight vs overweight
ol=60
ol=65
ol=27
ol=49
ol=65
ol=61
ol=24
ol=46
ol=8
ol=8
ol=11
ol=9
ol=20
ol=23
ol=7
Domain
12_vs_22 −a both
12_vs_22 −c male
0
10
20
30
−log10(FDR)
B
TCGA: GBM. cgc follow up tumor status. TUMOR FREE vs WITH TUMOR
oligodendrocyte development
oligodendrocyte differentiation
neuron part
paranode region of axon
ol=4
ol=5
ol=17
ol=3
0
0.5
1
1.5
−log10(FDR)
C
GTEx: Pituitary. BMI, body temperature, and hypertension
myelination
axon ensheathment
ensheathment of neurons
myelin sheath
structural constituent of myelin sheath
ol=4
ol=4
ol=4
ol=5
ol=4
Domain
BP
CC
MF
0
2
4
6
−log10(FDR)
D
TCGA: GBM. cgc follow up tumor status. TUMOR FREE vs WITH TUMOR: Oligodendrocyte pathway genes
CNP
LPAR1
CNTN2
NKX6-2
ASPA
TCGA: GBM. cgc follow up tumor status. TUMOR FREE vs WITH TUMOR: Neuronal pathway genes
MAP7
ERMN
LPAR1
GJC2
SH2D5
QDPR
TTLL7
NCDN
CNTN2
CPEB1
DBNDD2
SH3GL3
SPTBN4
LGI3
TPRN
RASGRF1
KNDC1
GTEx: Pituitary. BMI, body temperature, and hypertension: Oligodendrocyte pathway genes
DBNDD2
SH2D5
SPTBN4
GJC2
NCDN
TPRN
LGI3
CNTN2
QDPR
ERMN
MAP7
SH3GL3
TTLL7
LPAR1
KNDC1
RASGRF1
CPEB1
E
GTEx: Pituitary. MHASCITES Ascites. no vs yes
GTEx: Cells EBV−transformed lymphocytes. MHCANCERNM cancer non metastatic. no vs yes
GTEx: Skin Sun Exposed. MHASTHMA Asthma A. no vs yes
SRP030628 rc−brain: Dorsolateral prefrontal cortex. Infant vs Child stratified by male
SRP050892: H1 stem cell derived neural lines. Isotretinoin vs untreated control
SRP026256: Epithelial. transformed mesenchymal vs mesenchymal with OVOL1
SRP050892: H1 stem cell derived neural lines. Dioxin vs untreated control
TCGA: BRCA. xml er level cell percentage category. 60−69% vs 70−79%
SRP050892: H1 stem cell derived neural lines. Cytosine AZA−D−arabinofuranoside vs untreated control
TCGA: STAD. xml stage event pathologic stage. Stage IIIA vs Stage IIIC
TCGA: KIRC. xml neoplasm histologic grade. G2 vs G3
TCGA: STAD. xml stage event pathologic stage. Stage IIA vs Stage IIIA
SRP030628 rc−brain: Dorsolateral prefrontal cortex. Child vs Adolescent stratified by male
GTEx: Brain Cerebellar Hemisphere. MHDLYSIS Dialysis treatment. no vs yes
GTEx: Skin Sun Exposed. MHT2D type 2 diabetes. no vs yes
ol=68
ol=83
ol=57
ol=70
ol=81
ol=113
ol=56
ol=54
ol=90
ol=52
ol=89
ol=41
ol=20
ol=61
ol=72
Domain
12_vs_
22 −a
both
0
5
1
1
5
−log10(FD
F
SRP030628 rc−brain: Dorsolateral prefrontal cortex. Child vs Adolescent stratified by male
neuron differentiation
neurogenesis
generation of neurons
nervous system development
regulation: neuron differentiation
ol=24
ol=25
ol=24
ol=28
ol=14
Domain
BP
0
2
4
6
−log10(FDR)
G
VCAM1
MYCN
NR2F1
TUBB2B
GRIP1
DNMT3A
TOP2B
SOX11
CCDC88A
SOX4
KLF7
SHTN1
NKX2-2
RAC3
TRIM67
TNC
DPYSL3
STMN2
CASP3
DCX
DPYSL5
HES5
ISLR2
LRFN4

**Supplementary Figure S5. Differentially expressed genes (DEGs) between PD12 vs. PD22 male and female mice enriched against 3162 human differential gene expression comparisons using the Differential Expression Enrichment Tool (DEET) and ActivePathways.** A) Barplot of ActivePathways enrichment of upregulated DEGs integrating male and female comparisons. Rows are the top 15 most enriched DEG comparisons, and the X-axis is the FDR-adjusted p-value of enrichment. The number of overlapping DEGs between PD12 vs. PD22 mice and the human comparison is designated with the "ol=" in each comparison. The colour of each bar represents which sexes in the PD12 vs. PD22 comparison contributed to enrichment. B) Barplot displaying the top 5 most enriched gene ontology (GO) terms of the upregulated DEGs overlapping between the PD12 vs. PD22 comparisons and the "TCGA: GBM. cgc follow up tumour status TUMOR FREE vs WITH TUMOR" comparison in DEET. C) Barplot displaying the top 5 most enriched GO terms of the upregulated DEGs overlapping between the PD12 vs. PD22 comparisons and the hypothalamic-regulated comparisons in the pituitary gland. For B and C. Rows are different gene ontologies, and the X-axis is the FDR-adjusted p-value of enrichment. Bar colour designates whether the gene set is part of GO: biological process, molecular function, or cellular component. D) GeneMANIA plots of the overlapping genes between upregulated DEGs between PD12 vs PD22 comparisons and the Oligodendrocyte-related pathways in "TCGA: GBM. cgc follow up tumour status TUMOR FREE vs WITH TUMOR" (left), neuron-related pathways in "TCGA: GBM. cgc follow up tumour status TUMOR FREE vs WITH TUMOR" (center), and of the "hypothalamic-regulated comparisons in the pituitary gland". E) Barplot of ActivePathways enrichment of downregulated DEGs integrating male and female comparisons. Rows are the top 15 most enriched DEG comparisons and the X-axis is the FDR-adjusted p-value of enrichment. The number of overlapping DEGs between PD12 vs. PD22 mice and the human comparison is designated with the “ol=” in each comparison. F) Barplot displaying the top 5 most enriched GO terms of the downregulated DEGs overlapping between the PD12 vs. PD22 comparisons and the child vs adolescent comparison in the male PFC in humans. G) GeneMANIA plot of the overlapping genes between upregulated DEGs between PD12 vs PD22 comparisons and the child vs adolescent comparison in the male PFC in humans.

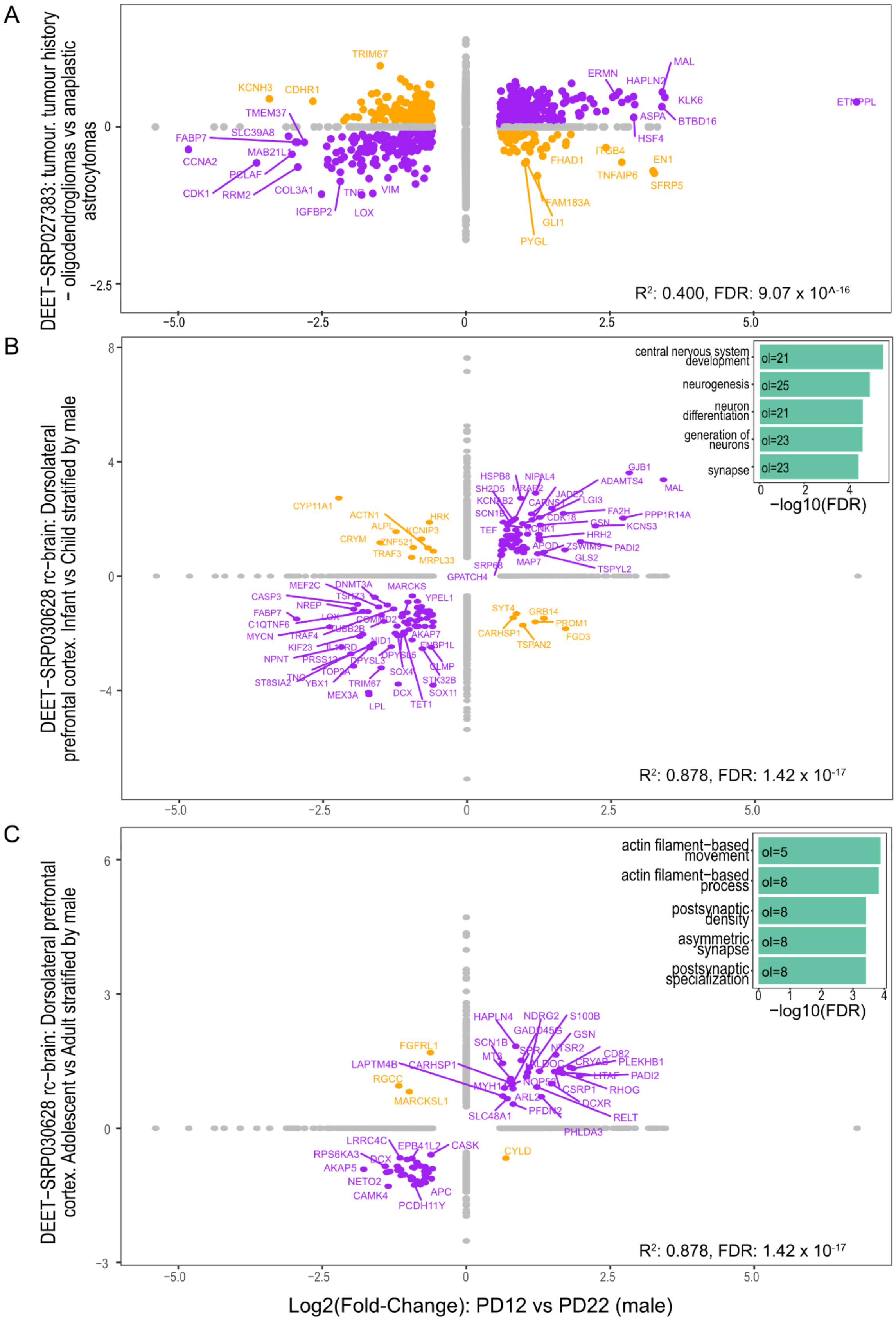

A
DEET−SRP027383: tumour. tumour history − oligodendrogliomas vs anaplastic astrocytomas
R²: 0.400, FDR: 9.07 x 10^-16
B
DEET−SRP030628 rc−brain: Dorsolateral prefrontal cortex. Infant vs Child stratified by male
central nervous system development
neurogenesis
neuron differentiation
generation of neurons
synapse
−log10(FDR)
R²: 0.878, FDR: 1.42 x 10-17
C
DEET−SRP030628 rc−brain: Dorsolateral prefrontal cortex. Adolescent vs Adult stratified by male
actin filament−based movement
actin filament−based process
postsynaptic density
asymmetric synapse
postsynaptic specialization
−log10(FDR)
R²: 0.878, FDR: 1.42 x 10-17
Log2(Fold-Change): PD12 vs PD22 (male)

**Supplementary Figure S6. Correlation plots between PD12 vs. PD22 males and the human comparisons within the DEET database that contain the top 3 most correlated DEGs.** A) correlation between the log2FC of PD12 vs. PD22 (male) and the log2FCs between tumour history - oligodendrogliomas vs. anaplastic astrocytomas. B) Correlation between the log2FC of PD12 vs. PD22 (male) and the log2FCs between infant vs child samples in the human pre-frontal cortex. B) Correlation between the log2FC of PD12 vs. PD22 (male) and the log2FCs between adolescent vs. adult samples in the human pre-frontal cortex. In B and C, the barplot represents the top 5 most enriched gene ontology (GO) terms of overlapping DEGs with the same sign of fold-change (i.e., purple points on the scatter plot). Rows are different GO terms, and the X-axis is the FDR-adjusted p-value of gene-set enrichment.

A

B

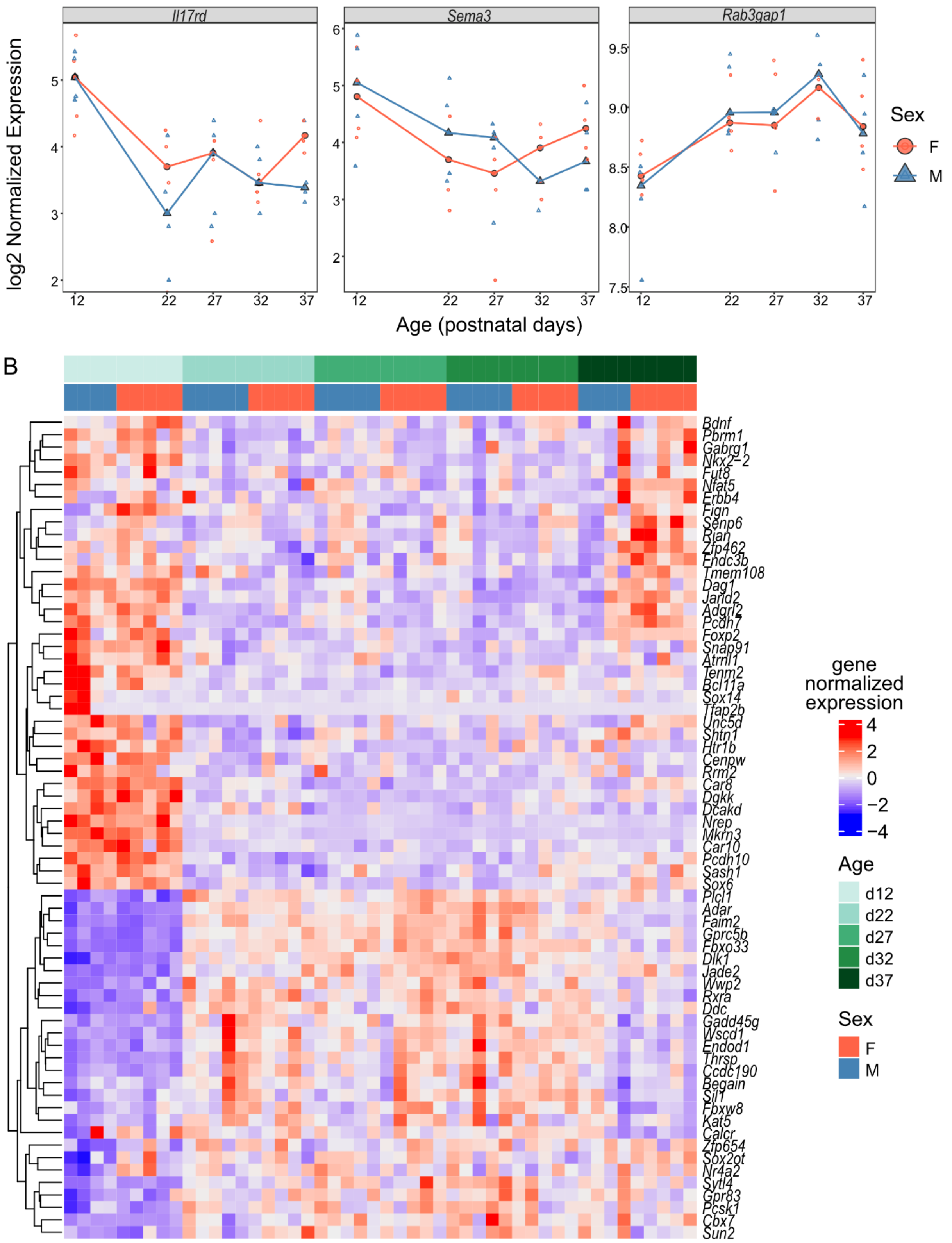

Il17rd
Sema3
Rab3gap1
log2 Normalized Expression
Age (postnatal days)
Sex
F
M
gene normalized expression
Age
d12
d22
d27
d32
d37

**Supplementary Figure S7. Summary of differentially expressed hypogonadotropic hypogonadism genes and puberty genome-wide association study (GWAS) genes.** A) Distribution of normalized counts from hypogonadotropic hypogonadism genes. The X-axis is age, and the Y-axis is log2-transformed RUVseq and ERCC-spike in normalized counts. Red lines and circles represent female samples, while blue lines and triangles represent male samples. B) Gene expression heatmap of all UTR-seq samples. Rows are DEGs that have been identified as puberty GWAS genes, and columns are samples. The heatmap is populated by log2 RUV-seq normalized gene expression profiles. Columns are annotated by age and sex and labeled by replicate number.

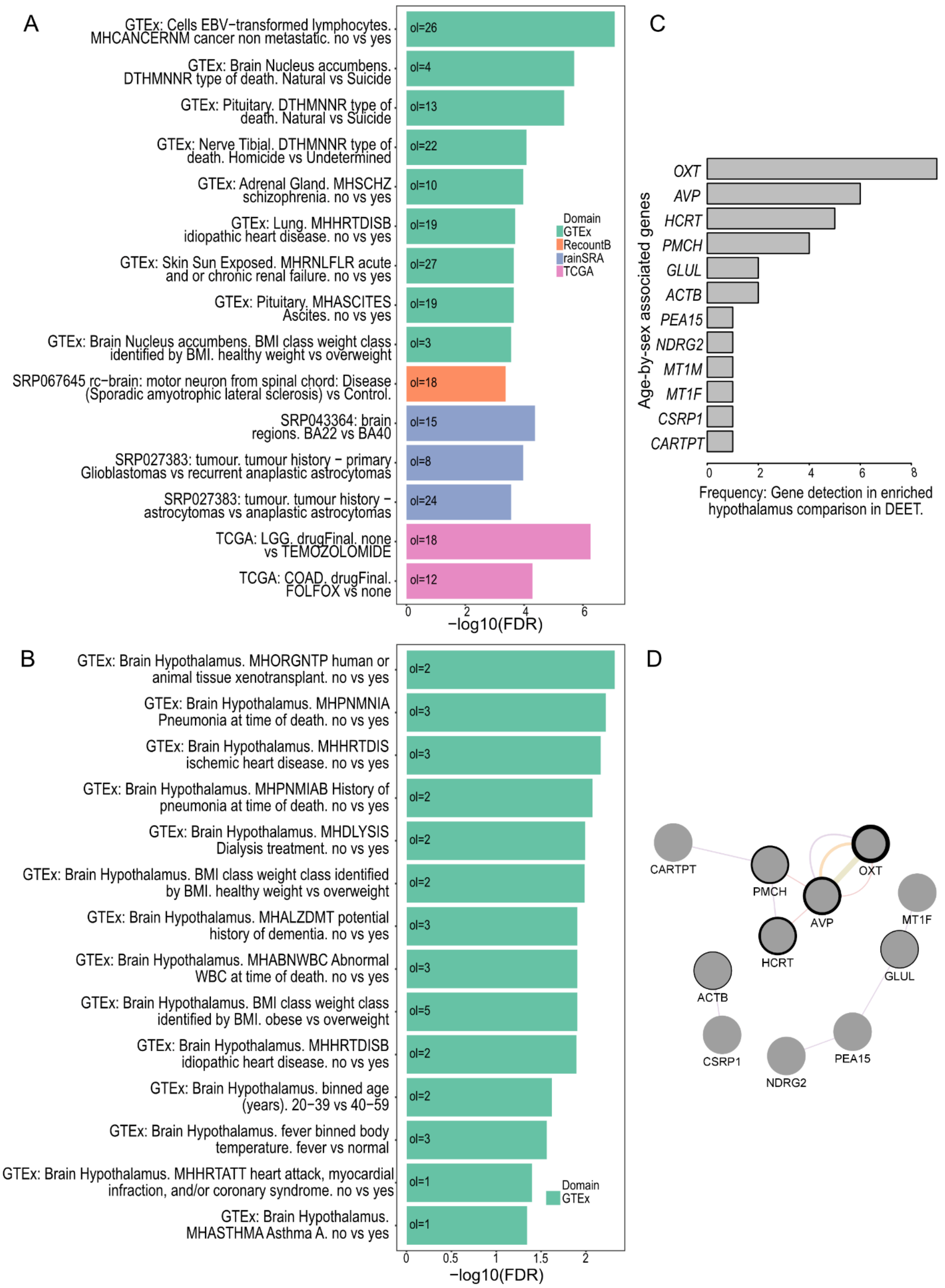
A
GTEx: Cells EBV−transformed lymphocytes. MHCANCERNM cancer non metastatic. no vs yes
GTEx: Brain Nucleus accumbens. DTHMNNR type of death. Natural vs Suicide
GTEx: Pituitary. DTHMNNR type of death. Natural vs Suicide
GTEx: Nerve Tibial. DTHMNNR type of death. Homicide vs Undetermined
GTEx: Adrenal Gland. MHSCHZ schizophrenia. no vs yes
GTEx: Lung. MHHRTDISB idiopathic heart disease. no vs yes
GTEx: Skin Sun Exposed. MHRNLFLR acute and or chronic renal failure. no vs yes
GTEx: Pituitary. MHASCITES Ascites. no vs yes
GTEx: Brain Nucleus accumbens. BMI class weight class identified by BMI. healthy weight vs overweight
SRP067645 rc−brain: motor neuron from spinal chord: Disease (Sporadic amyotrophic lateral sclerosis) vs Control.
SRP043364: brain regions. BA22 vs BA40
SRP027383: tumour. tumour history − primary Glioblastomas vs recurrent anaplastic astrocytomas
SRP027383: tumour. tumour history − astrocytomas vs anaplastic astrocytomas
TCGA: LGG. drugFinal. none vs TEMOZOLOMIDE
TCGA: COAD. drugFinal. FOLFOX vs none
ol=26
ol=4
ol=13
ol=22
ol=10
ol=19
ol=27
ol=19
ol=3
ol=18
ol=15
ol=8
ol=24
ol=18
ol=12
Domain
GTEx
RecountB
rainSRA
TCGA
0
2
4
6
−log10(FDR)
B
GTEx: Brain Hypothalamus. MHORGNTP human or animal tissue xenotransplant. no vs yes
GTEx: Brain Hypothalamus. MHPNMNIA Pneumonia at time of death. no vs yes
GTEx: Brain Hypothalamus. MHHRTDIS ischemic heart disease. no vs yes
GTEx: Brain Hypothalamus. MHPNMIAB History of pneumonia at time of death. no vs yes
GTEx: Brain Hypothalamus. MHDLYSIS Dialysis treatment. no vs yes
GTEx: Brain Hypothalamus. BMI class weight class identified by BMI. healthy weight vs overweight
GTEx: Brain Hypothalamus. MHALZDMT potential history of dementia. no vs yes
GTEx: Brain Hypothalamus. MHABNWBC Abnormal WBC at time of death. no vs yes
GTEx: Brain Hypothalamus. BMI class weight class identified by BMI. obese vs overweight
GTEx: Brain Hypothalamus. MHHRTDISB idiopathic heart disease. no vs yes
GTEx: Brain Hypothalamus. binned age (years). 20−39 vs 40−59
GTEx: Brain Hypothalamus. fever binned body temperature. fever vs normal
GTEx: Brain Hypothalamus. MHHRTATT heart attack, myocardial infraction, and/or coronary syndrome. no vs yes
GTEx: Brain Hypothalamus. MHASTHMA Asthma A. no vs yes
ol=2
ol=3
ol=3
ol=2
ol=2
ol=2
ol=3
ol=3
ol=5
ol=2
ol=2
ol=3
ol=1
ol=1
Domain
GTEx
0
0.5
1
1.5
2
−log10(FDR)
C
Age-by-sex associated genes
OXT
AVP
HCRT
PMCH
GLUL
ACTB
PEA15
NDRG2
MT1M
MT1F
CSRP1
CARTPT
0
2
4
6
8
Frequency: Gene detection in enriched hypothalamus comparison in DEET.
D
CARTPT
PMCH
OXT
AVP
MT1F
HCRT
GLUL
ACTB
CSRP1
PEA15
NDRG2

**Supplementary Figure S8. Enrichment of age-by-sex associated genes against 3162 human RNA-seq comparisons stored in the DEET dataset.** Barplot of the top 15 most associated DE comparisons identified using the "DEET_enrich" tool. B) Barplot to the top (up-to) 15 most associated DE comparisons occurring within "Hypothalamus" tissue in DEET. Rows are different comparisons, and the X-axis represents the degree of enrichment for each comparison. Bar colours represent the dataset class where the comparison was sourced from (i.e., recount-brain, TCGA, GTEx, SRA). C) Barplot displaying the frequency of hypothalamus comparisons that age-by-sex associated genes were detected as DE within. D) GeneMANIA of the genes in C). Nodes are genes, and edge-width is the number of hypothalamus comparisons where the gene is DE. Edges represent connections between genes stored within the GeneMANIA dataset. Purple edges represent co-expression, red edges represent protein-protein interactions, orange edges represent predicted interactions, and yellow-green edges represent shared protein domains.

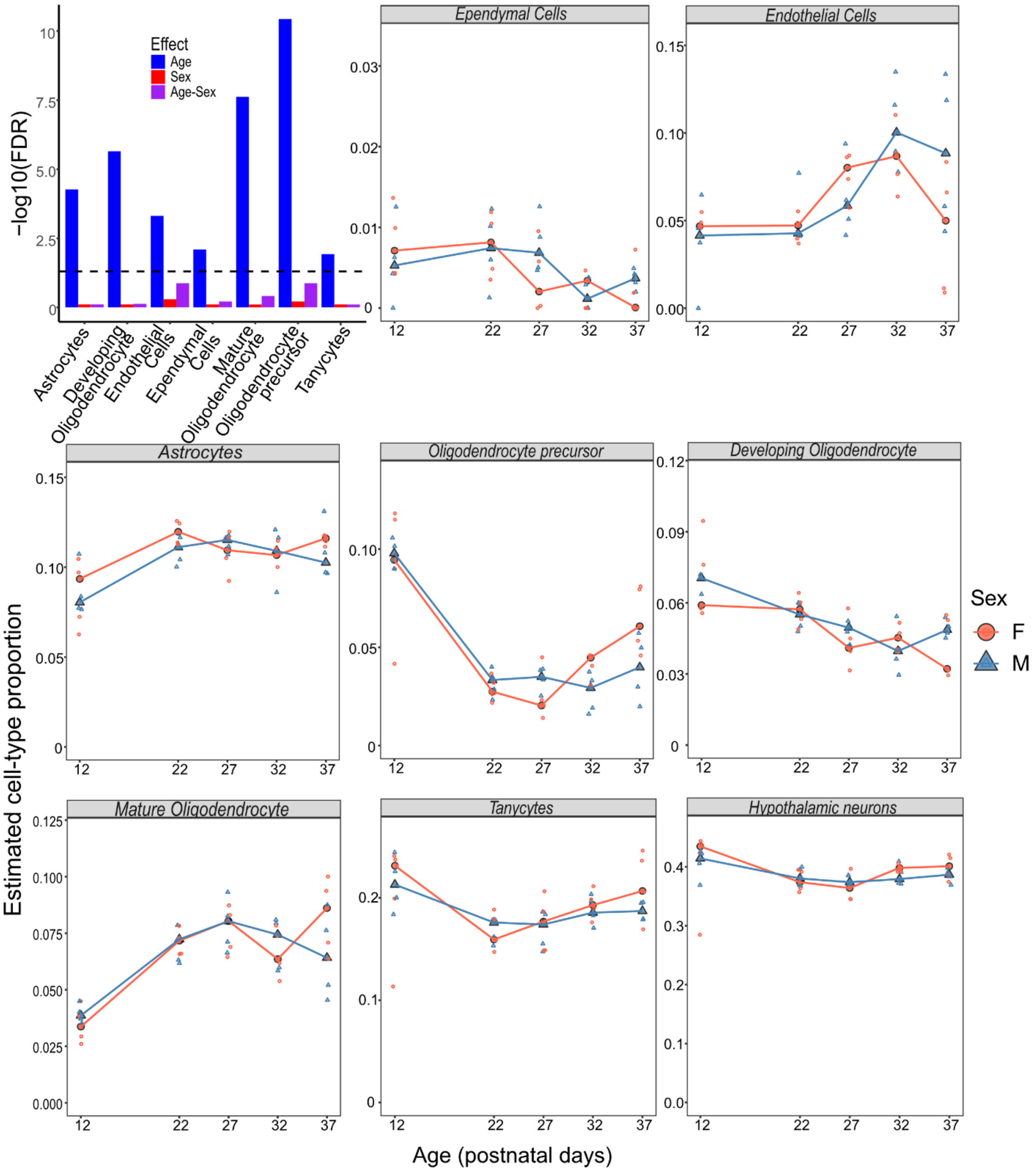


**Supplementary Figure S9. Distribution of cell-type proportions measured from MuSiC-NNLS in cell-types that were predicted to have >3% of the total sample.** The top-leftmost plot displays cell-type proportion association between age, sex, and an age-by-sex interaction across puberty. The X-axis is different cell-types, and Y-axis is the -log10(FDR-adjusted p-value) of a two-way ANOVA. The remaining plots are the cell-type distributions of each cell-type across ages and sex. The X-axis is age, and the Y-axis is the estimated cell-type proportion. Red lines and circles represent female samples, while blue lines and triangles represent male samples.

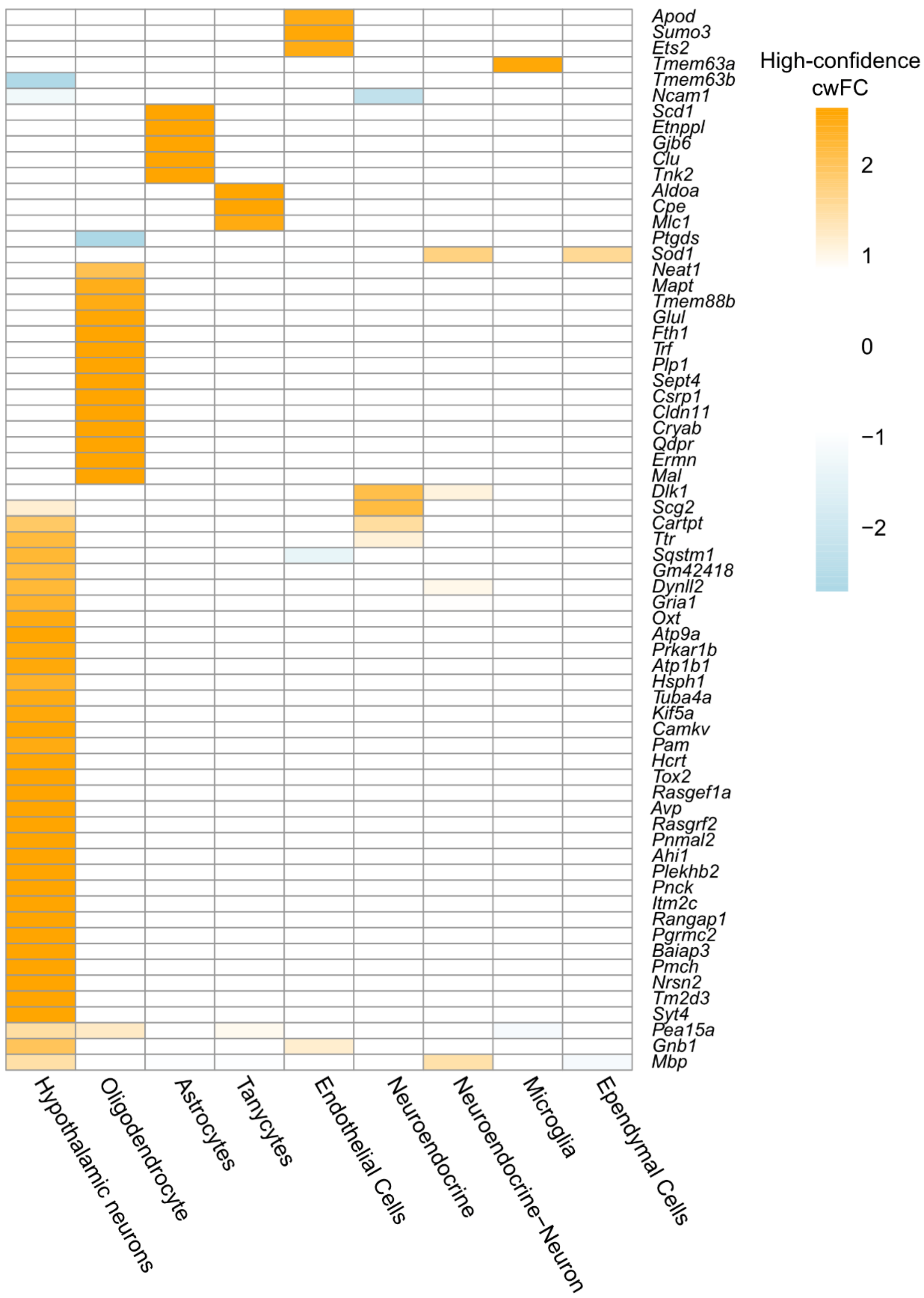
High-confidence cwFC
2
1
0
−1
−2
Apod
Sumo3
Ets2
Tmem63a
Tmem63b
Ncam1
Scd1
Etnppl
Gjb6
Clu
Tnk2
Aldoa
Cpe
Mlc1
Ptgds
Sod1
Neat1
Mapt
Tmem88b
Glul
Fth1
Trf
Plp1
Sept4
Csrp1
Cldn11
Cryab
Qdpr
Ermn
Mal
Dlk1
Scg2
Cartpt
Ttr
Sqstm1
Gm42418
Dynll2
Gria1
Oxt
Atp9a
Prkar1b
Atp1b1
Hsph1
Tuba4a
Kif5a
Camkv
Pam
Hcrt
Tox2
Rasgef1a
Avp
Rasgrf2
Pnmal2
Ahi1
Plekhb2
Pnck
Itm2c
Rangap1
Pgrmc2
Baiap3
Pmch
Nrsn2
Tm2d3
Syt4
Pea15a
Gnb1
Mbp
Hypothalamic neurons
Oligodendrocyte
Astrocytes
Tanycytes
Endothelial Cells
Neuroendocrine
Neuroendocrine−Neuron
Microglia
Ependymal Cells

**Supplementary Figure S10. Heatmap of gene-normalized cell-weighted fold-changes (cwFold-changes) of the 129 age-by-sex associated genes and are DE in the complementary direction in the scRNA-seq data.** Rows are genes, and columns are cell-types.

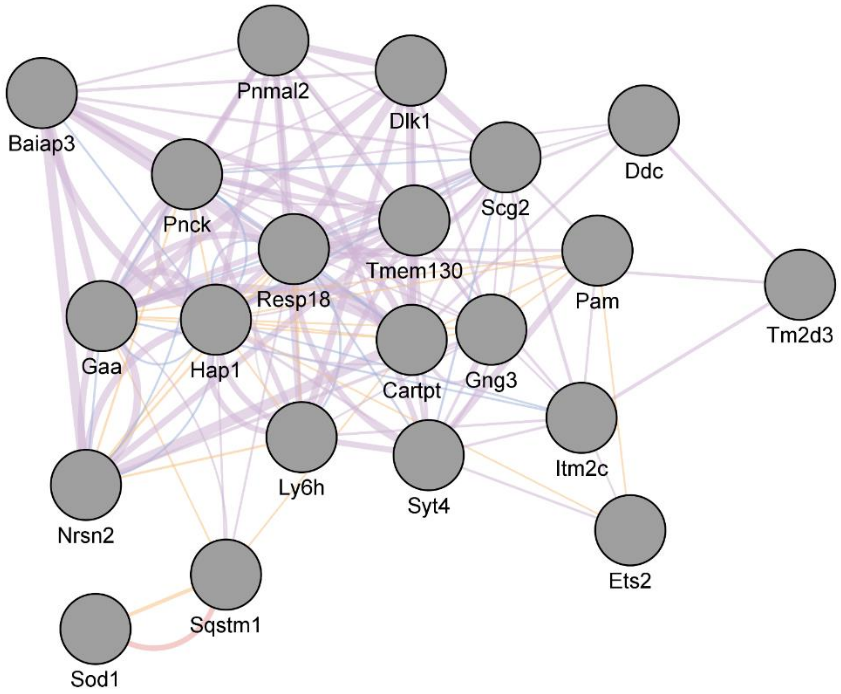


**Supplementary Figure S11. GeneMANIA plot of the 21 neuron- neuroendocrine-mapping age-by-sex associated genes that are detected as translated in hypothalamic LepRb+ neurons in Trap-seq from Allison et al., 2015.** Nodes are genes, and edges are connections between genes pre-computed within the GeneMANIA database. Purple edges are co-expression, blue edges are co-localization, orange edges are predicted interactions, and red edges are validated physical interactions.

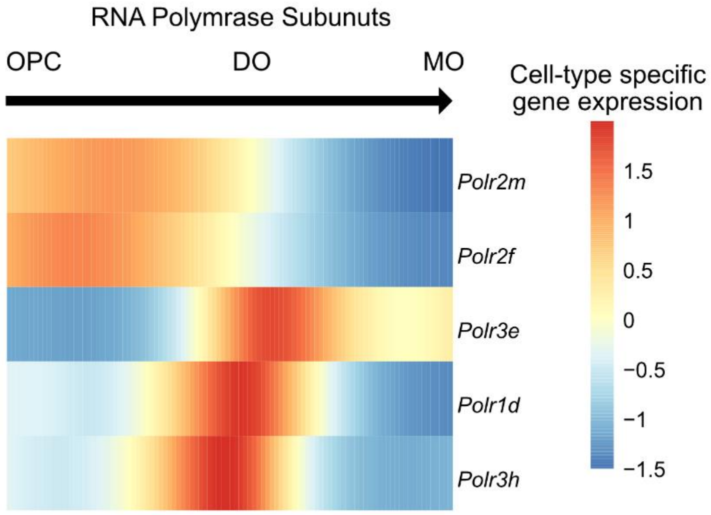


**Supplementary Figure S12. Pseudotime of hypothalamic oligodendrocyte development. Heatmap of RNA polymerase subunit genes associated with pseudotime.** For a gene to be included, it must be associated with an age-by-sex interaction (i.e., varimax 16), mapping to oligodendrocyte precursor cells, developing oligodendrocytes, or mature oligodendrocytes with scMappR, and associate with pseudotime. For B-D, rows are genes associated with pseudotime. Columns are portions of the pseudotime trajectory blocked into 200 smoothers using tradeSeq. Heat is measured by scaling the predicted smoothers with the scale function in R.

### Supplementary Tables

**Supplementary Table S1**. Summary of RNA-seq sample quality and read mapping using Qualimap.

| Sample | secondary.alignments | total.alignments | ambiguous.alignments | no.feature.assigned | not.aligned | intronic | intergenic | overlapping.exon |
|---|---|---|---|---|---|---|---|---|
| WL1000_hypothal_mmus_Hd12F4 | 897512 | 4018830 | 80655 | 763433 | 0 | 78865 (22.28%) | 184568 (7.1%) | 49499 (1.91%) |
| WL1001_hypothal_mmus_Hd12F5 | 816150 | 3585337 | 69658 | 643259 | 0 | 83482 (21.12%) | 159777 (6.98%) | 40399 (1.76%) |
| WL1002_hypothal_mmus_Hd12F6 | 762491 | 3457266 | 72863 | 580768 | 0 | 34638 (19.46%) | 146130 (6.54%) | 44583 (2%) |
| WL1003_hypothal_mmus_Hd12F7 | 279824 | 1247718 | 25700 | 219815 | 0 | 63879 (20.49%) | 55936 (6.99%) | 15521 (1.94%) |
| WL1004_hypothal_mmus_Hd22M1 | 307948 | 1394625 | 27980 | 249736 | 0 | 84828 (20.52%) | 64908 (7.2%) | 16658 (1.85%) |
| WL1005_hypothal_mmus_Hd22M3 | 592869 | 2739735 | 54882 | 501622 | 0 | 78294 (21.12%) | 123328 (6.89%) | 34390 (1.92%) |
| WL1006_hypothal_mmus_Hd22M4 | 2720538 | 12528767 | 248130 | 2201778 | 0 | 644000 (20.13%) | 557778 (6.83%) | 153833 (1.88%) |
| WL1007_hypothal_mmus_Hd22M5 | 881128 | 4383875 | 87658 | 912494 | 0 | 11960 (24.02%) | 200534 (6.77%) | 60085 (2.03%) |
| WL1008_hypothal_mmus_Hd22M6 | 690060 | 3258466 | 65121 | 647344 | 0 | 93449 (22.89%) | 153895 (7.14%) | 45699 (2.12%) |
| WL1009_hypothal_mmus_Hd22F2 | 759355 | 3581985 | 70305 | 722470 | 0 | 57977 (23.53%) | 164493 (6.94%) | 45677 (1.93%) |
| WL1010_hypothal_mmus_Hd22F3 | 1647709 | 7618744 | 154872 | 1333203 | 0 | 92209 (19.95%) | 340994 (6.86%) | 93922 (1.89%) |
| WL1011_hypothal_mmus_Hd22F4 | 364287 | 1660806 | 33918 | 297505 | 0 | 18103 (20.22%) | 79402 (7.36%) | 21834 (2.02%) |
| WL1012_hypothal_mmus_Hd22F5 | 435274 | 2028310 | 42008 | 373957 | 0 | 83348 (21.35%) | 90609 (6.83%) | 24995 (1.88%) |
| WL1013_hypothal_mmus_Hd22F6 | 165422 | 743356 | 14826 | 128262 | 0 | 3753 (19.65%) | 34509 (7.23%) | 9612 (2.01%) |
| WL1014_hypothal_mmus_Hd27M2 | 750596 | 3463314 | 70971 | 597842 | 0 | 47068 (19.81%) | 150774 (6.68%) | 41963 (1.86%) |
| WL1015_hypothal_mmus_Hd27M3 | 5268708 | 25387546 | 474237 | 5437437 | 0 | 210552 (24.81%) | 1226885 (7.23%) | 305102 (1.8%) |
| WL1016_hypothal_mmus_Hd27M4 | 824809 | 3874816 | 73567 | 809951 | 0 | 31837 (24.64%) | 178114 (6.95%) | 45122 (1.76%) |
| WL1017_hypothal_mmus_Hd27M5 | 1236481 | 5874076 | 112991 | 1179686 | 0 | 96444 (23.02%) | 283242 (7.27%) | 73924 (1.9%) |
| WL1018_hypothal_mmus_Hd27M6 | 1390603 | 6244161 | 129394 | 1017383 | 0 | 47621 (18.59%) | 269762 (6.71%) | 85312 (2.12%) |
| WL1019_hypothal_mmus_Hd27F2 | 391261 | 1723825 | 33907 | 300226 | 0 | 16944 (19.8%) | 83282 (7.6%) | 20266 (1.85%) |
| WL1020_hypothal_mmus_Hd27F3 | 390777 | 1791860 | 37798 | 315971 | 0 | 35247 (20.2%) | 80724 (6.93%) | 25460 (2.19%) |
| WL1021_hypothal_mmus_Hd27F4 | 744720 | 3444402 | 71345 | 587431 | 0 | 37615 (19.49%) | 149816 (6.67%) | 44706 (1.99%) |
| WL1022_hypothal_mmus_Hd27F5 | 757263 | 3548183 | 73741 | 614409 | 0 | 60079 (19.79%) | 154330 (6.64%) | 41129 (1.77%) |
| WL1023_hypothal_mmus_Hd27F6 | 1013292 | 4858922 | 94491 | 1036959 | 0 | 03315 (24.75%) | 233644 (7.2%) | 62796 (1.93%) |
| WL1024_hypothal_mmus_Hd32M1 | 1142682 | 5354579 | 104587 | 1026548 | 0 | 94014 (22.47%) | 232534 (6.58%) | 64028 (1.81%) |
| WL1025_hypothal_mmus_Hd32M3 | 802247 | 3639021 | 69649 | 729937 | 0 | 54140 (23.4%) | 175797 (7.42%) | 46421 (1.96%) |
| WL1026_hypothal_mmus_Hd32M4 | 3858259 | 18278342 | 400304 | 3055582 | 0 | 257573 (18.76%) | 798009 (6.63%) | 281030 (2.34%) |
| WL1027_hypothal_mmus_Hd32M5 | 2276859 | 10379096 | 212262 | 1674474 | 0 | 235536 (18.4%) | 438938 (6.54%) | 111605 (1.66%) |
| WL1028_hypothal_mmus_Hd32M6 | 478545 | 2165257 | 44653 | 365020 | 0 | 65072 (18.99%) | 99948 (7.16%) | 29703 (2.13%) |
| WL1029_hypothal_mmus_Hd32F1 | 1079679 | 5094237 | 108694 | 901223 | 0 | 82194 (20.36%) | 219029 (6.54%) | 59979 (1.79%) |
| WL1030_hypothal_mmus_Hd32F3 | 881626 | 4027460 | 81429 | 756465 | 0 | 66635 (21.63%) | 189830 (7.24%) | 53837 (2.05%) |
| WL1031_hypothal_mmus_Hd32F4 | 2151473 | 10256559 | 191158 | 2270076 | 0 | 786957 (26.12%) | 483119 (7.06%) | 134149 (1.96%) |
| WL1032_hypothal_mmus_Hd32F5 | 977740 | 4555696 | 86541 | 972881 | 0 | 61766 (25.36%) | 211115 (7.03%) | 53947 (1.8%) |
| WL1033_hypothal_mmus_Hd32F6 | 961263 | 4458001 | 85422 | 955138 | 0 | 51860 (25.61%) | 203278 (6.93%) | 57490 (1.96%) |
| WL1034_hypothal_mmus_Hd37M1 | 1179887 | 5368416 | 113994 | 878820 | 0 | 49877 (18.7%) | 228943 (6.59%) | 66941 (1.93%) |
| WL1035_hypothal_mmus_Hd37M2 | 894074 | 4001554 | 85646 | 650723 | 0 | 79049 (18.68%) | 171674 (6.69%) | 48683 (1.9%) |
| WL1036_hypothal_mmus_Hd37M4 | 957481 | 4542704 | 85859 | 974106 | 0 | 48473 (24.95%) | 225633 (7.52%) | 57455 (1.91%) |
| WL1037_hypothal_mmus_Hd37M6 | 766851 | 3605096 | 67089 | 711797 | 0 | 43101 (22.86%) | 168696 (7.1%) | 38265 (1.61%) |
| WL1038_hypothal_mmus_Hd37F1 | 728635 | 3672249 | 66250 | 935061 | 0 | 61037 (30.29%) | 174024 (6.93%) | 47627 (1.9%) |
| WL1039_hypothal_mmus_Hd37F2 | 724558 | 3706350 | 67763 | 938410 | 0 | 59058 (29.81%) | 179352 (7.04%) | 48804 (1.92%) |
| WL1040_hypothal_mmus_Hd37F3 | 350644 | 1673101 | 32111 | 369817 | 0 | 87075 (25.85%) | 82742 (7.45%) | 21102 (1.9%) |
| WL1041_hypothal_mmus_Hd37F4 | 1037322 | 5087204 | 93475 | 1270935 | 0 | 027029 (29.83%) | 243906 (7.08%) | 70500 (2.05%) |
| WL1042_hypothal_mmus_Hd37F6 | 2209519 | 10306322 | 195179 | 2037279 | 0 | 570309 (23.19%) | 466970 (6.9%) | 126087 (1.86%) |
| WL994_hypothal_mmus_Hd12M1 | 1244301 | 5619012 | 117675 | 929960 | 0 | 10424 (19.56%) | 219536 (6.04%) | 71688 (1.97%) |
| WL996_hypothal_mmus_Hd12M4 | 334884 | 1541831 | 31885 | 281893 | 0 | 15910 (21.46%) | 65983 (6.56%) | 19792 (1.97%) |
| WL997_hypothal_mmus_Hd12M5 | 1127055 | 5144960 | 105470 | 952741 | 0 | 32288 (21.87%) | 220453 (6.58%) | 64239 (1.92%) |
| WL998_hypothal_mmus_Hd12M6 | 786105 | 3532137 | 73342 | 605942 | 0 | 51613 (19.88%) | 154329 (6.79%) | 43983 (1.94%) |
| WL999_hypothal_mmus_Hd12F2 | 649604 | 2927720 | 58216 | 573712 | 0 | 43754 (23.39%) | 129958 (6.85%) | 35694 (1.88%) |

**Supplementary Table S2**. Correlation and number of cell-types detected (>1% of the total population) between cell-type proportions from RNA-seq deconvolution and cell-type proportions from scRNA-seq data in the mouse hypothalamus.

| Deconvolution Approach | PD12 vs PD14 (scRNA-seq): $R^2$ | PD37 vs PD45 (scRNA-seq): $R^2$ | All samples vs. PD14 and PD45: $R^2$ | Detected cell-types (>1% of population) |
|---|---|---|---|---|
| MuSiC | 0.445 | 0.202 | 0.271 | 8 |
| NNLS | 0.743 | 0.638 | 0.696 | 7 |
| CPM | -0.334 | -0.396 | -0.416 | 11 |
| Cibersort | -0.374 | -0.359 | -0.413 | 6 |
| Cibersortx | -0.356 | -0.381 | -0.414 | 6 |
| WGCNA | 0.194 | 0.328 | 0.308 | 5 |
| DCQ | 0.310 | 0.504 | 0.486 | 4 |
| DeconRNAseq | 0.499 | 0.420 | 0.438 | 11 |
| BayesPrism | -0.084 | 0.162 | 0.187 | 6 |
